# A conserved genetic interaction between Spt6 and Set2 regulates H3K36 methylation

**DOI:** 10.1101/364521

**Authors:** Rajaraman Gopalakrishnan, Fred Winston

**Author notes:** Corresponding author: Fred Winston.

## Abstract

The transcription elongation factor Spt6 and the H3K36 methyltransferase Set2 are both required for H3K36 methylation and transcriptional fidelity in *Saccharomyces cerevisiae*. By selecting for suppressors of a transcriptional defect in an *spt6* mutant, we have isolated dominant *SET2* mutations (*SET2^sup^* mutations) in a region encoding a proposed autoinhibitory domain. The *SET2^sup^* mutations suppress the H3K36 methylation defect in the *spt6* mutant, as well as in other mutants that impair H3K36 methylation. ChIP-seq studies demonstrate that the H3K36 methylation defect in the *spt6* mutant, as well as its suppression by a *SET2^sup^* mutation, occur at a step following the recruitment of Set2 to chromatin. Other experiments show that a similar genetic relationship between Spt6 and Set2 exists in *Schizosaccharomyces pombe*. Taken together, our results suggest a conserved mechanism by which the Set2 autoinhibitory domain requires multiple interactions to ensure that H3K36 methylation occurs specifically on actively transcribed chromatin.

## Introduction

The histone chaperone Spt6 is a highly conserved transcription elongation factor required for many aspects of transcription and chromatin structure. Spt6 binds directly to Rpb1, the largest subunit of RNA polymerase II (RNAPII)^1–6^, to histones and nucleosomes^7–9^, and to the essential transcription factor Spn1/Iws1^9–12^. Mutations in *S. cerevisiae SPT6* cause genome-wide changes in histone occupancy^13–16^ and impair several histone modifications, including H3K36 di- and trimethylation (H3K36me2/me3) catalyzed by the H3K36 methyltransferase Set2^17–20^. Mutations in *SPT6* also cause greatly elevated levels of transcripts that arise from within coding regions on both sense and antisense strands, known as intragenic transcription^15,21–24^. Intragenic transcription has recently emerged as a mechanism to express alternative genetic information within a coding region (for example, ^25–29^).

Regulation of intragenic transcription by Spt6 occurs, at least in part, by its regulation of H3K36 methylation, as a deletion of *SET2* also causes genome-wide expression of intragenic transcripts^17,30,31^. Set2 normally represses intragenic transcription via its association with RNAPII during transcription elongation, resulting in H3K36me2/me3 over gene bodies^32–34^. This histone modification is required for the co-transcriptional function of the Rpd3S histone deacetylase complex^17,35–38^. Deacetylation by Rpd3S over transcribed regions is believed to maintain a repressive environment that prevents intragenic transcription. Regulation of intragenic transcription by H3K36 methylation is conserved as depletion of *SETD2* (a human orthologue of yeast *SET2*) also results in the genome-wide expression of intragenic transcripts^39^.

Set2-dependent H3K36me2/me3 is regulated by several factors in addition to Spt6. These include members of the PAF complex^33,40^, as well as the Rpb1 CTD kinases Ctk1^33,41^ and Bur1^18,40^. Furthermore, there is strong evidence that a nucleosomal surface composed of specific residues of histones H2A, H3, and H4 near the entry and exit point of nucleosomal DNA form a substrate recognition surface for Set2^42,43^. The H3 N-terminal tail itself has also been shown to be required for Set2 activity and mutant analysis suggests that intra-tail interactions^44^ and *cis-trans* isomerization of the N-terminal H3 tail^45^ control Set2 activity. The combined influence of all of these factors shows that Set2 activity is highly regulated to ensure that it occurs co-transcriptionally on a chromatin template.

Multiple domains within Set2 regulate its catalytic activity in order to ensure that it functions during transcription elongation. The C-terminal region of Set2 contains the Set2-Rpb1 interacting domain (SRI domain) which interacts with the Ser2- and Ser5-phosphorylated carboxy-terminal domain (CTD) of Rpb1^46^ and which binds nucleosomal DNA^47^. A deletion of the SRI domain causes loss of H3K36 methylation^19^. In addition, a nine amino acid sequence in the N-terminal region of Set2 mediates the interaction of Set2 with histone H4 and this domain is also required for Set2 catalytic activity^43^. The central region of Set2 has been characterized as an autoinhibitory domain^47^, as deletions throughout this region result in increased H3K36 methylation^47^. However, the functional role of this domain is unknown.

The initial goal of our study was to identify factors that regulate Spt6-mediated intragenic transcription. To do this, we carried out a selection for suppressor mutations that inhibit intragenic transcription in an *spt6* mutant, where intragenic transcripts are widespread^22,23^. We identified 20 independent, dominant mutations in *SET2* (*SET2^sup^* mutations) that encode a cluster of amino acid changes in the Set2 autoinhibitory domain. The isolation of these mutants led us to study the function of the autoinhibitory domain *in vivo*. Our results show that our *SET2^sup^* mutations suppress H3K36me2/me3 defects in *spt6* and other transcription elongation factor mutants, as well as in *set2* mutants that normally abolish Set2 activity. In addition, we show that the loss of H3K36me2/me3 in *spt6-1004* and its suppression by the *SET2^sup^* mutations both occur genome-wide, primarily at a step beyond Set2 recruitment. Finally we show that orthologous *SET2^sup^* mutations in *S. pombe* also partially rescue the H3K36 methylation defect in an *S. pombe spt6* mutant. Taken together, our results have revealed new insights into the regulation of Set2 and suggest that the autoinhibitory domain monitors multiple Set2 interactions that are required for its function *in vivo*.

## Results

### Isolation and analysis of dominant *SET2* mutations that suppress intragenic transcription in an *spt6-1004* mutant

To identify factors that regulate intragenic transcription, we selected for mutations that suppress this class of transcription in an *spt6-1004* mutant^21^, which allows extensive intragenic transcription^22,23^. To select for suppressors, we constructed two reporters using characterized intragenic transcription start sites in the *FLO8*^21^ and *STE11*^48^ genes (Fig. 1a; Methods). In the *FLO8-URA3* reporter, intragenic transcription confers sensitivity to 5-FOA, while in the *STE11-CAN1* reporter, intragenic transcription confers sensitivity to canavanine. To select for mutations that suppress intragenic transcription, we constructed *spt6-1004* strains that contained both reporters and selected for resistance to both 5-FOA and canavanine (5-FOA^R^ Can^R^). The double selection reduced the likelihood of isolating *cis*-acting mutations in either reporter, thereby enriching for mutants that generally affect intragenic transcription.

**Fig. 1.**
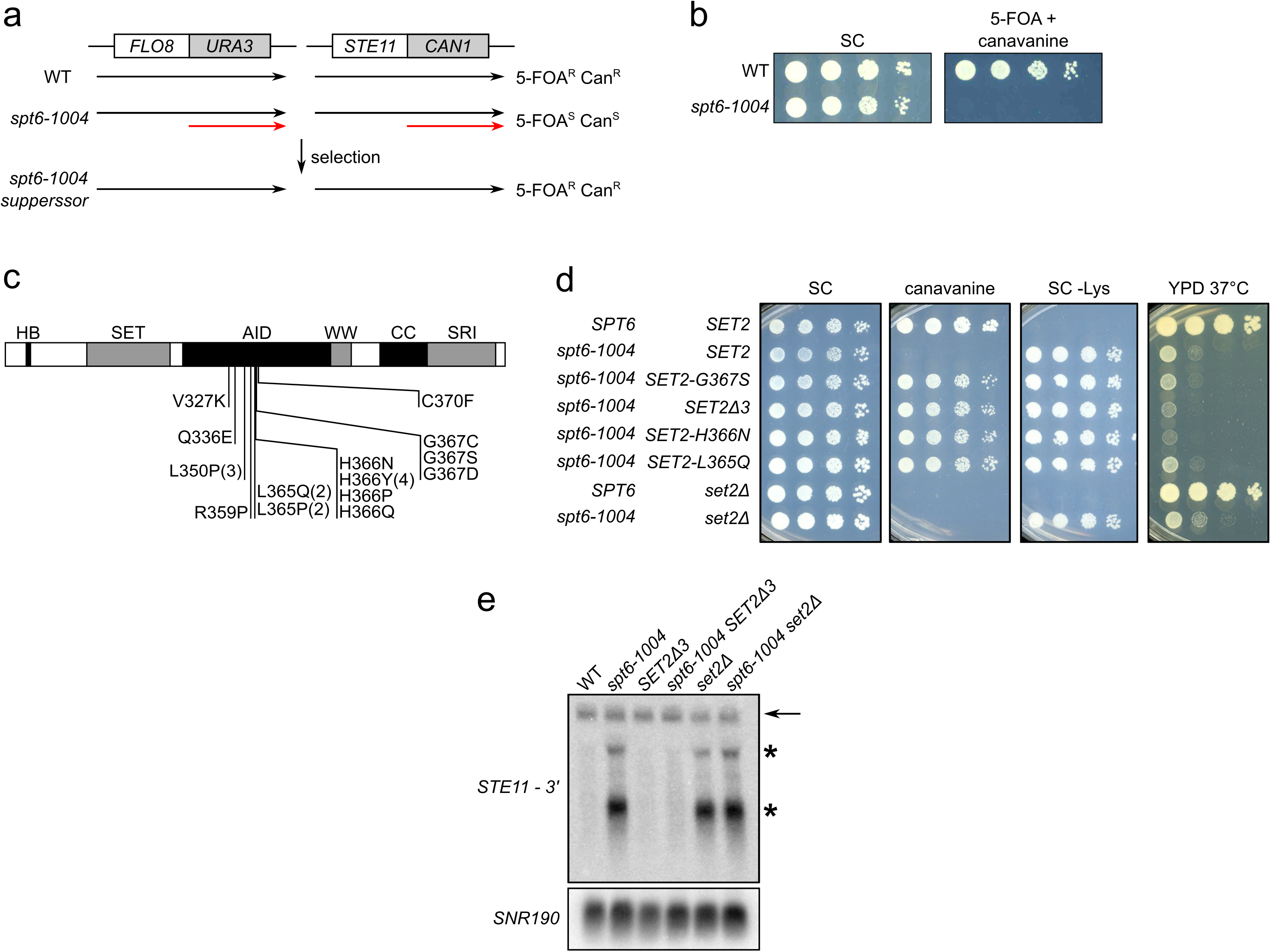
Isolation of mutations that suppress intragenic transcription in *spt6-1004*. **a**, Shown at the top are diagrams of the two reporters and the selection used to isolate suppressor mutations. Shown below each are the transcripts made in wild-type (WT), *spt6-1004*, and *spt6-1004* strains with a suppressor. In *spt6-1004* mutants, the *FLO8-URA3* reporter confers 5-FOA sensitivity and the *STE11-CAN1* reporter confers canavanine sensitivity due to expression of the intragenic transcripts, shown in red. **b**, Spot tests of cells grown at 34°C that show the selective conditions for the mutant selection. **c**, A diagram of the Set2 protein that depicts the amino acid changes caused by the dominant *SET2^sup^* mutations. The numbers in parentheses indicate the number of times the same mutation was isolated if more than once. One isolate had two point mutations that coded for L365P and H366Y in the same protein. The letters above the rectangle indicate the identified domains in Set2: HB, histone binding domain; SET, the catalytic domain; AID, the autoinhibitory domain; the WW domain; CC, a coiled coil motif; and SRI, the Set2 Rpb1 interacting domain. **d**, Spot tests of cells grown at 34°C to assay suppression of intragenic transcription in *spt6-1004* by SET2^*sup*^ mutations using the *STE11-CAN1* reporter. **e**, Northern analysis of the *STE11* gene, using a probe from the 3’ region of *STE11*, to assay suppression of intragenic transcription in *spt6-1004* by a *SET2^sup^* mutation. In wild type there is a single full-length *STE11* transcript (denoted by the arrow), while in the *spt6-1004* mutant, there are two intragenic transcripts (denoted by the asterisks) in addition to the full-length transcript. *SNR190* served as the loading control.

We isolated and characterized 20 independent mutants. By standard genetic tests, we showed that all 20 mutations were dominant. We then tested eight mutants by crosses and showed that the 5-FOA^R^ Can^R^ phenotype was caused by a single mutation in each strain and that the mutations were tightly linked to each other, with no recombinants found in any of seven crosses (10 tetrads/cross). To identify candidate mutations, we performed whole genome sequencing of these eight suppressor strains and identified single base pair changes in the *SET2* gene in all eight mutants, suggesting that these are the causative mutations that suppress intragenic transcription. Sequencing of the *SET2* gene in the other 12 suppressors also revealed mutations in *SET2*. The 20 mutations (Table S1) are clustered within a small region of *SET2* encoding a previously-identified autoinhibitory domain^47^.

To verify that the dominant *SET2* mutations are causative for suppression of intragenic transcription in *spt6-1004*, we recreated three of the identified *SET2^sup^* mutations in the *spt6-1004* parental reporter strains. As 13 of the 20 *SET2^sup^* mutations are within three adjacent codons (365-367) (Fig. 1c), we decided to test three of these mutations, *SET2-L365Q*, *SET2-H366N*, and *SET2-G367S*, and a fourth mutation (*SET2Δ3*) that deleted these three *SET2* codons. In all four cases, the reconstructed mutants were 5-FOA^R^ and Can^R^, showing that each of the *SET2* mutations was causative (Fig. 1d). Suppression was specific for intragenic transcription as the mutants still had other *spt6* mutant phenotypes, including Spt^−^ and temperature-sensitive growth (Fig. 1d). Deletion of the entire *SET2* gene does not suppress intragenic transcription in an *spt6-1004* background (Fig. 1d), demonstrating that our *SET2^sup^* mutations do not cause loss of Set2 activity. To assay the effect of a *SET2^sup^* mutation on levels of an intragenic transcript, we performed Northern blots, looking at *STE11* transcripts, using a strain with a wild-type *STE11* gene. Our results showed that the *SET2Δ3* mutation strongly suppressed *STE11* intragenic transcript levels in an *spt6-1004* mutant, to levels similar to that in wild-type cells (Fig. 1e). Suppression by the *SET2Δ3* mutation suggests that the suppression phenotype occurs by impairment of the Set2 autoinhibitory domain. Taken together, our results show that mutations that change or remove amino acids in the Set2 autoinhibitory domain suppress intragenic transcription in an *spt6-1004* mutant.

### *SET2^sup^* mutations rescue H3K36 di- and trimethylation in an *spt6-1004* mutant

Given that all of our suppressor mutations were in *SET2*, we tested whether they suppress the H3K36me2/me3 defect in *spt6-1004*, using quantitative Western blots. Compared to the *spt6-1004* single mutant, where H3K36me3 is undetectable, our results show that four different *SET2^sup^ spt6-1004* double mutants have a substantial level of H3K36me3, approximately 10-40% of the level of a wild-type strain (Fig. 2a,b). In addition, all of the other originally isolated *SET2^sup^* mutants, tested once, restored H3K36me3 to varying extents in an *spt6-1004* background (data not shown). Furthermore, we constructed a series of short deletions that removed segments of the Set2 autoinhibitory domain and found that they also suppressed the H3K36me2/me3 defect in *spt6-1004* to a similar degree as the *SET2^sup^* mutations (Supplementary Fig. 1). Thus, multiple types of changes in the Set2 autoinhibitory domain partially bypass the requirement of Set2 for Spt6.

**Fig. 2.**
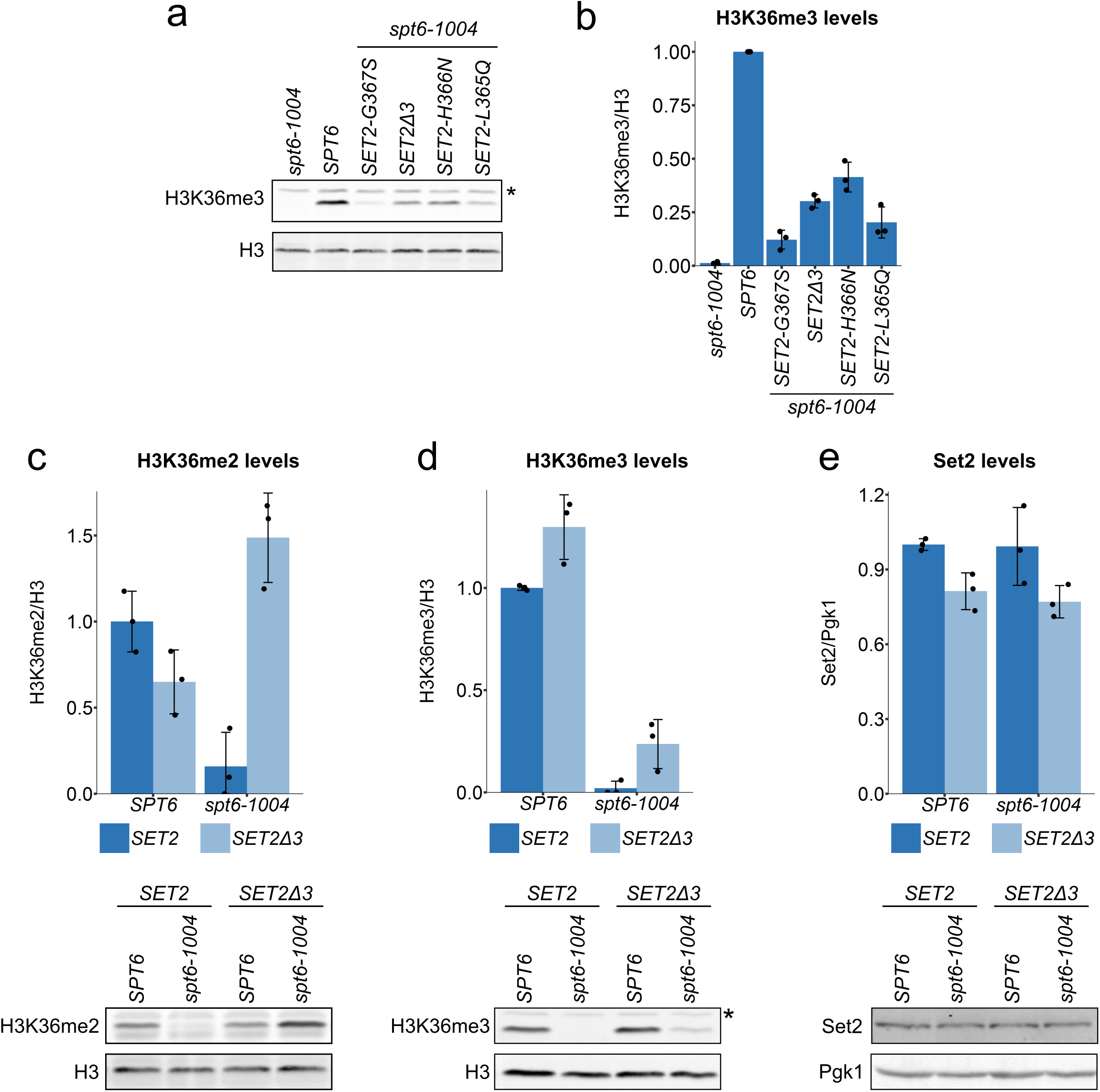
*SET2Δ3* suppresses the H3K36 methylation defect in *spt6-1004*. **a**, A Western blot showing the levels of H3K36me3 and total histone H3 in four different *SET2^sup^* mutants. The asterisk denotes a non-specific band. **b**, Quantification of H3K36me3 relative levels of the strains shown in (a). **c,d,e**, Quantification of Western blots assaying the levels of H3K36me2 (**c**), H3K36me3 (**d**) and Set2 (**e**) in wild-type and *spt6-1004* strains with or without the *SET2Δ3* mutation, normalized to their respective loading controls. For all bar graphs, the black dots represent the individual data points for three experiments and the bars show the mean +/- standard deviation. A representative Western blot is shown below each bar graph.

To determine the effect of a *SET2^sup^* mutation in an otherwise wild-type background, we compared the effects of the *SET2Δ3* mutation on H3K36me2 and H3K36me3 levels with and without an *spt6-1004* mutation. In an *spt6-1004 SET2Δ3* double mutant, H3K36me3 levels are approximately 25% of wild-type levels and H3K36me2 levels are approximately 150% of wild-type (Fig. 2c,d). In the *SET2Δ3* single mutant, there appears to be hyperactivation of Set2 activity, as we observed greater levels of H3K36me3 and slightly decreased levels of H3K36me2 compared to wild type (Fig. 2c,d). Importantly, these changes in H3K36 methylation are not caused by elevated levels of Set2 protein (Fig. 2e). Taken together, our results suggest that the Set2 autoinhibitory domain makes Set2 activity dependent upon Spt6.

To test whether *SET2^sup^* mutations can also suppress depletion of the Spt6 protein in addition to suppressing the *spt6-1004* mutation, we conditionally depleted Spt6 via an auxin-inducible degron^49^. In a wild-type *SET2* background, as expected, we observed decreased levels of H3K36me2/me3 upon Spt6 depletion (Supplementary Fig. 2). Set2 levels also decreased during the time course of this experiment, although this occurs later than the loss of H3K36 methylation. When Spt6 is depleted in the *SET2Δ3* background, we observed increased levels of H3K36me2/me3 during the depletion compared to the wild-type *SET2* background, although the levels eventually decreased as Set2 protein levels decreased. Despite the decreasing levels of Set2, these results show that *SET2^sup^* mutations partially bypass the H3K36me2/me3 defects caused by depletion of Spt6.

### *SET2^sup^* mutations suppress *spt6-1004* via the Set2/Rpd3S pathway

We also performed two sets of experiments to verify that the *SET2^sup^* mutations function via H3K36 methylation and the function of Rpd3S. First, to confirm that the *SET2^sup^* mutations exert their phenotype by restoring methylation of H3K36 rather than by some other event, we compared *spt6-1004 SET2-H366N* strains that express either wild-type histone H3 or an H3K36A mutant. Our results showed that the *spt6-1004 SET2-H366N* strain expressing H3K36A was no longer able to suppress intragenic transcription (Fig. 3a); therefore, H3K36 methylation is necessary for suppression of intragenic transcription. Second, as H3K36me2/me3 is required for the function of the Rpd3S histone deacetylase complex^17,35–38^, we assayed whether suppression of *spt6-1004* required a functional Rpd3S complex, by testing a strain lacking the Rpd3S component Rco1. Our results showed that *rco1Δ* reversed the suppression phenotype, similar to the H3K36A mutant (Fig. 3b), showing that functional Rpd3S is necessary for suppression of intragenic transcription by *SET2^sup^* mutations. Together, these results demonstrate that methylation at H3K36 and the subsequent activation of Rpd3S confers suppression by the *SET2^sup^* mutations.

**Fig. 3.**
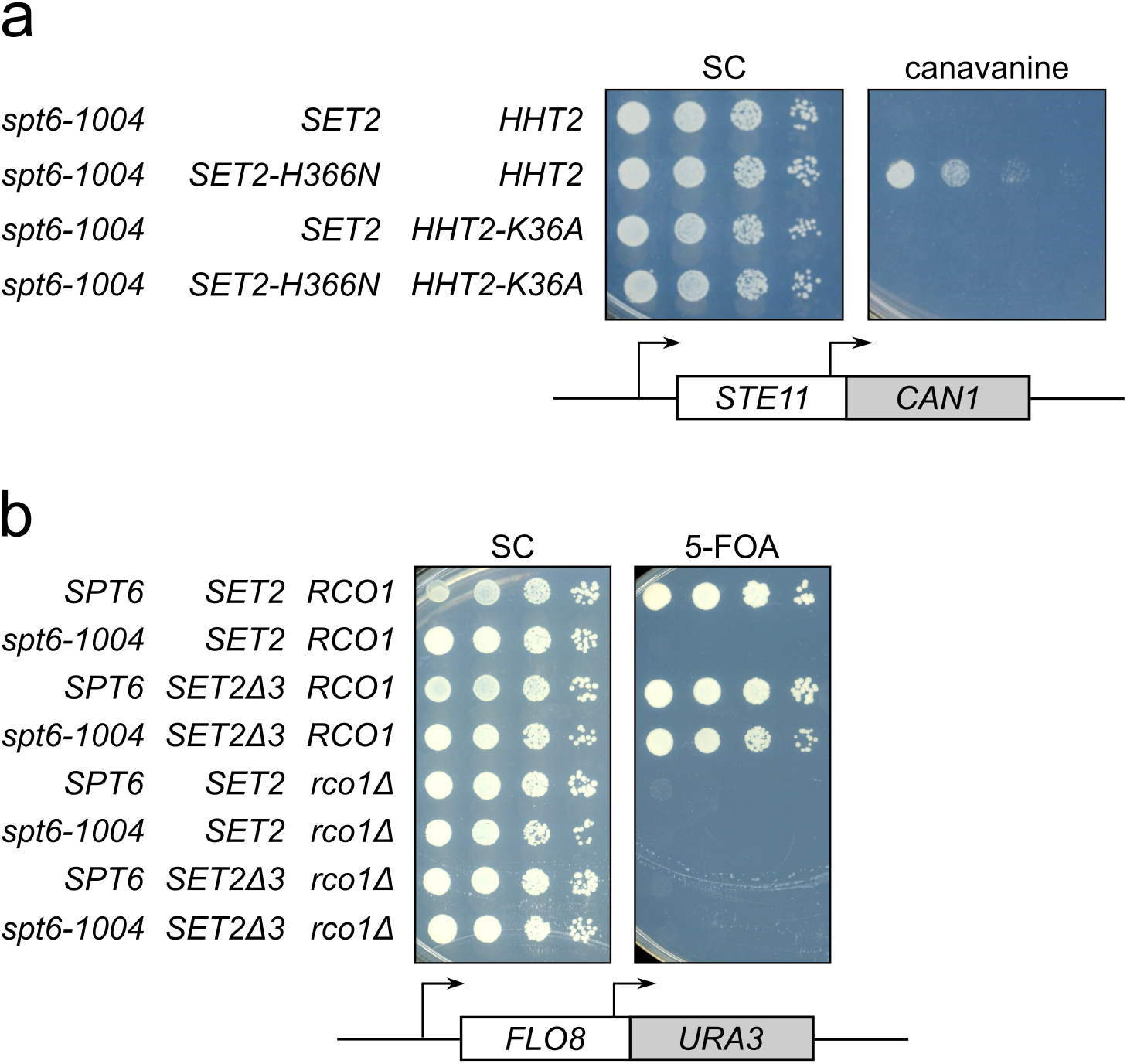
*SET2^sup^* mutations suppress *spt6-1004* via the Set2/Rpd3S pathway. **a**, Spot tests of cells grown at 34°C assaying expression of the *STE11-CAN1* reporter, showing the effect of the H3K36A mutation on the suppression phenotype. **b**, Spot tests of cells grown at 34°C, assaying the effect of *rco1Δ* on the expression of the *FLO8-URA3* reporter in suppressor strains.

### *SET2^sup^* mutations suppress H3K36 methylation defects that occur in other transcription elongation factor mutants

We wanted to test whether *SET2^sup^* mutations can suppress the loss of other functions that are required for both H3K36 methylation and repression of intragenic transcription. In particular, we tested the PAF complex and Ctk1 which, along with Spt6, have been proposed to be part of a feed-forward mechanism that regulates transcription elongation^50^. For the PAF complex, we tested *paf1Δ* and *ctr9Δ*, both of which cause loss of H3K36me3, with no detectable effect on either H3K36me2 or Set2 protein levels (Fig. 4a-c;^40^). *SET2Δ3* strongly suppressed the H3K36me3 defect of both *paf1Δ* and *ctr9Δ* (Fig. 4b). In a *ctk1Δ* mutant, there are decreased Set2 protein levels and loss of both H3K36me3 and H3K36me2 (Fig. 4b;^19,41,50^. *SET2Δ3* strongly suppressed the H3K36me2 defect in *ctk1Δ*, restoring it to a level greater than in wild-type strains (Fig. 4b), although it had no effect on H3K36me3 or on the diminished Set2 protein levels (Fig. 4a,c). We also tested whether *SET2Δ3* suppresses intragenic transcription in these mutants. Our results showed that *SET2Δ3* suppressed intragenic transcription in *paf1Δ* and *ctr9Δ* mutants, but not in the *ctk1Δ* mutant (Fig. 4d). The latter result suggests that restoration of H3K36me2 but not H3K36me3 in *ctk1Δ* is insufficient for the repression of intragenic transcription. The bypass of the requirements for multiple factors by *SET2^sup^* mutations suggests that the Set2 autoinhibitory domain confers dependence upon these factors for Set2 function.

**Fig. 4.**
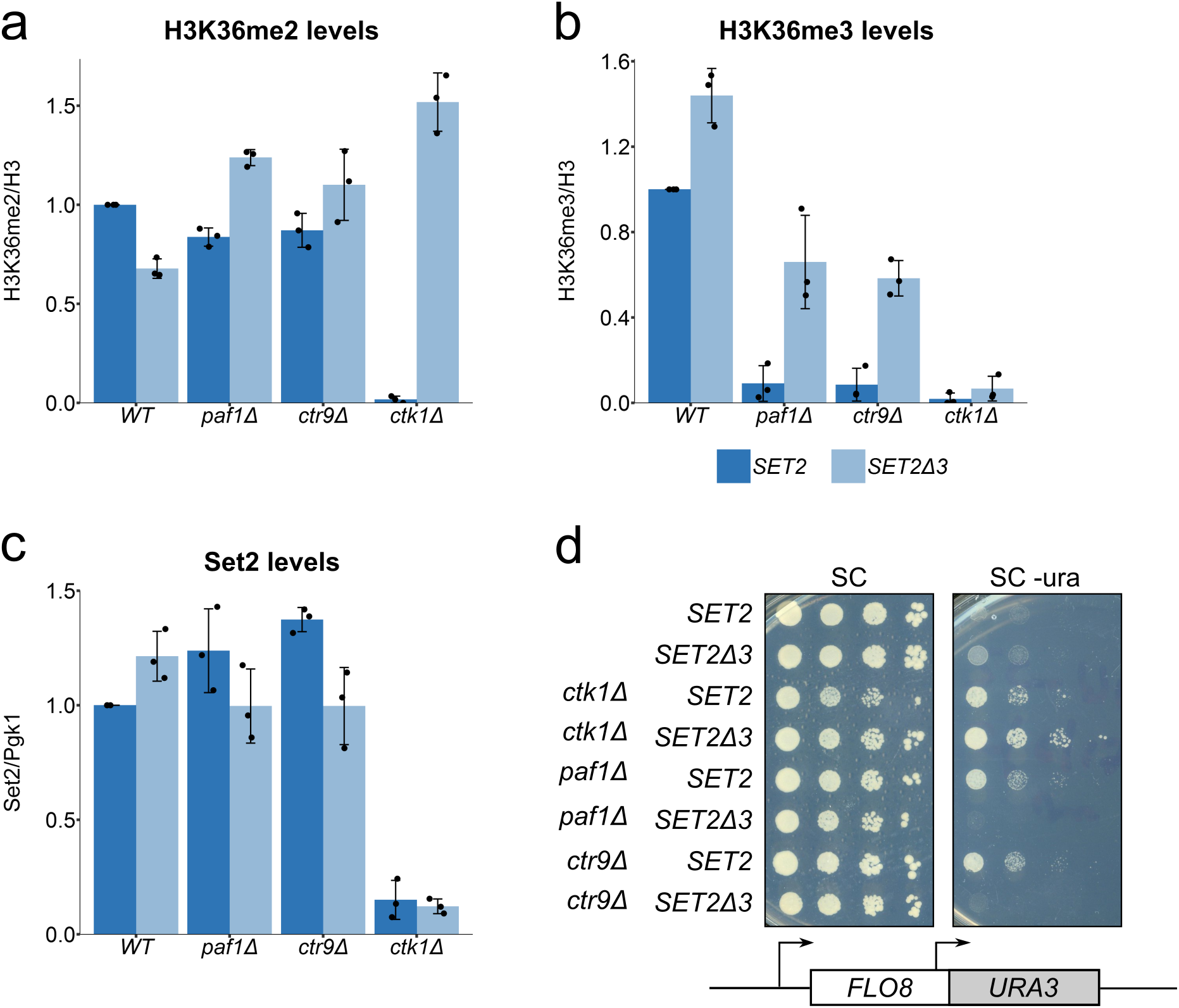
*SET2Δ3* suppress the H3K36me2/me3 defect in *ctk1Δ, paf1Δ and ctr9Δ*. **a,b,c**, Quantification of Western blots assaying H3K36me2 (**a**), H3K36me3 (**b**) and Set2 (**c**) levels in the indicated mutants with or without the *SET2Δ3* mutation, normalized to their respective loading controls. The black dots represent the individual data points for three experiments and the bars show the mean +/- standard deviation. **d**, Spot tests of cells grown at 30°C, assaying the effect of the *SET2Δ3* mutation on expression of the *FLO8-URA3* reporter in the indicated mutants. Since intragenic transcription is weak in these mutants, growth has been assayed on SC-Ura medium (see Methods).

### *SET2^sup^* mutations suppress the loss of Set2 domains normally required for its catalytic activity

We also investigated whether *SET2^sup^* mutations suppress the loss of two Set2 regulatory domains required for Set2 activity: the SRI domain, which binds to the RNAPII CTD^19,46^, and the HB domain, required for interaction with histones H2A and H4^43^. To do this we deleted portions of the *SET2* gene to remove one or both domains in the Set2 protein in a wild-type *SET2* gene and a *SET2Δ3* mutant (Fig. 5a). We then tested the new mutants for levels of H3K36me2, H3K36me3, and Set2. Our results showed that *SET2Δ3* suppresses both the *set2ΔSRI* and *set2ΔHB* mutations with respect to their H3K36 methylation defects (Fig. 5a,b, Supplementary Fig. 3). However, *SET2Δ3* is unable to rescue H3K36 methylation in a *set2ΔHB,ΔSRI* double mutant. This is not due to either altered recruitment or level of the mutant protein (Fig. 5b, Supplementary Fig. 3c). Consistent with the H3K36 methylation levels, *SET2Δ3* was able to suppress intragenic transcription in a *set2ΔSRI* but not in a *set2ΔHB,ΔSRI* strain (Fig. 5c). Our results suggest that the Set2 autoinhibitory domain monitors the interactions of the Set2 SRI and HB domains with RNAPII and nucleosomes, respectively.

**Fig. 5.**
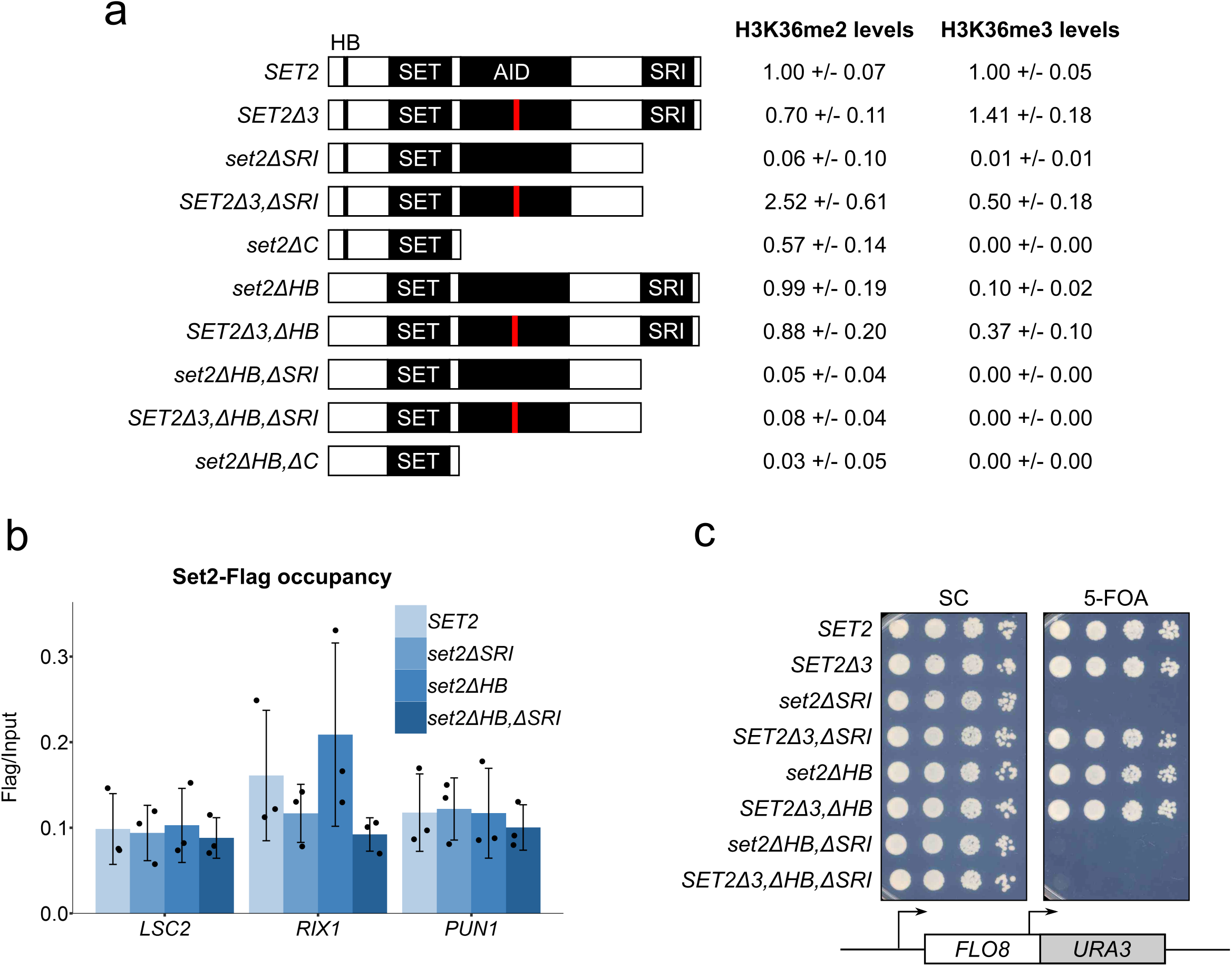
*SET2Δ3* rescues H3K36 methylation in *set2* mutants lacking the SRI or HB domains. **a**, The schematic depicts Set2 and a set of mutants missing the indicated domains, testing each for H3K36me2 and H3K36me3 levels by western analysis. The red line in the AID represents the position of the *SET2Δ3* mutation. The numbers next to each mutation show the levels of H3K36me2 and H3K36me3 in each mutant strain relative to a wild-type control (mean +/- standard deviation of at least three experiments). **b**, ChIP-qPCR assaying localization of FLAG-tagged Set2 wild-type and mutant proteins at the *LSC2*, *RIX1*, and *PUN1* genes. The black dots represent the individual data points for three experiments and the bars show the mean +/- standard deviation. ChIP-qPCR data has been spike-in normalized to the *S. pombe ACT1* gene. **c.** Spot tests of cells grown at 37°C, assaying expression of the *FLO8-URA3* reported in the indicated strains.

### The regulation of Set2 activity by Spt6 and Set2^sup^ mutants occurs at a step after the recruitment of Set2 to chromatin

Although the H3K36 methylation defect in *spt6-1004* mutants was discovered several years ago, there is little understanding of why Spt6 is required for this histone modification. Our *spt6-1004* strains, when grown at 30°C, have almost normal Set2 levels (Fig. 2d; Supplementary Fig. 4c); however, H3K36me2 or H3K36me3 are undetectable. Therefore, the requirement for Spt6 must be at a step other than regulation of Set2 stability. Two other possible mechanisms for regulation include the recruitment of Set2 to chromatin or the regulation of Set2 activity after its recruitment. To distinguish between these possibilities, as well as to better understand the suppression of *spt6-1004* by *SET2^sup^* mutations, we performed ChIP-seq for Set2-HA, Rpb1, H3K36me3, H3K36me2, and total H3. These experiments were performed in four genetic backgrounds: wild type, *spt6-1004*, *spt6-1004 SET2-H366N*, and *SET2-H366N*. Each condition was performed in duplicate and was highly reproducible (Supplementary Fig. 44a). We chose *SET2-H366N* as it was the strongest suppressor of the *spt6-1004* H3K36me3 defect. To permit quantitative comparisons of ChIP signals between different samples, we used *S. pombe* chromatin for spike-in normalization (Methods, Supplementary Fig. 4b).

Our results revealed new information regarding the H3K36 methylation defect caused by *spt6-1004* as well as the suppression of this defect by *SET2-H366N*. First, there was a large decrease genome-wide in H3K36me2 and H3K36me3 association with chromatin as compared to the wild-type strain (Fig. 6a,b), a result consistent with the observation that H3K36me2 and H3K36me3 are undetectable by westerns in *spt6-1004*. Second, in contrast to the large decrease in H3K36me2/me3, there was little decrease in the level of Set2 protein recruited across transcribed regions (Fig. 6c) when normalized to the level of Rpb1 (Supplementary Fig.5). Given these results, the defect in H3K36me2/me3 in the *spt6-1004* mutant must occur primarily at a level subsequent to Set2 recruitment to chromatin. Third, in the *spt6-1004 SET2-H366N* double mutant, we saw a genome-wide rescue of H3K36me2/me3 (Fig. 6a,b). Compared to the wild-type strain, this strain had generally increased levels of H3K36me2 and decreased levels of H3K36me3. Set2 localization to transcribed regions was similar to that in *spt6-1004* (Fig. 6c), showing that suppression by *SET2-H366N* was not due to increased recruitment to chromatin. Finally, the *SPT6 SET2-H366N* single mutant showed increased H3K36me3 and decreased H3K36me2 levels genome-wide as compared to wild type (Fig. 6a,b). However, in this strain the recruitment of Set2 to chromatin is modestly increased (Fig. 6c), which may be due to the higher level of the Set2-H366N-Flag protein (Supplementary Fig. 4c). No global changes in histone H3 occupancy were observed in any of the strains (Supplementary Fig. 5c,f). ChIP-qPCR results at individual genes were consistent with our ChIP-seq results (see *STE11* and *RIX1*, Fig. 6d-f; Supplementary Fig. 5d-f). In summary, our results show that Spt6 is required for Set2 function after its recruitment to chromatin and suggest that this requirement is dependent upon the Set2 autoinhibitory domain.

**Fig. 6.**
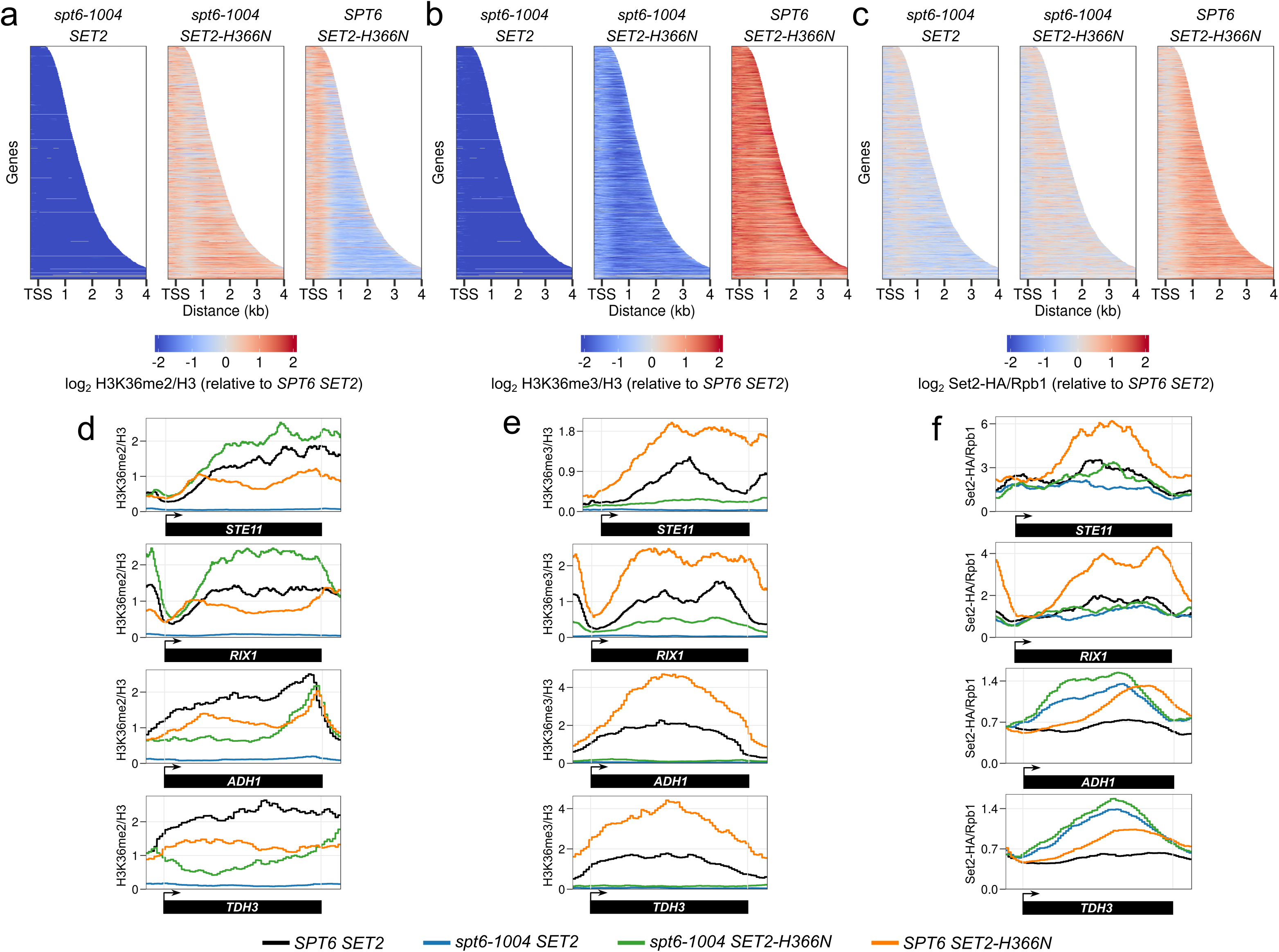
*SET2Δ3* rescues H3K36 methylation genome-wide in *spt6-1004*. **a, b, c** Heatmaps depicting H3K36me2 levels (relative to H3) (**a**), H3K36me3 levels (relative to H3) (**b**) and Set2-HA levels (relative to Rpb1) (**c**) for all non-overlapping genes that code for protein (n = 3522). All values that had a log fold change of below −2 or above 2 have been set to −2 or 2, respectively. **d.e.f**, H3K36me2 (**c**), H3K36me3 (**d**), and Set2-HA **(e)** levels at genes that show rescue of H3K36 methylation (*STE11*, *RIX1*) and genes where rescue of H3K36 methylation is not observed (*ADH1*, *TDH3*). All data has been normalized to an *S. pombe* spike-in control.

Although the effects we observed occurred at most genes, a set of 48 genes behaved differently. In contrast to most genes, which had increased levels of H3K36me2 and partial rescue of H3K36me3 in the *spt6-1004 SET2-H366N* strains as compared to wild type, this set of genes had reduced levels of H3K36me2 compared to wild type. These genes had slightly decreased occupancy of histone H3 as compared to wild type, but that decrease could not account for the decreased level of H3K36me2. In addition, Set2 recruitment was not impaired at these genes. Examples of two such genes, *ADH1* and *TDH3*, are shown in Fig. 6d-f. GO term analysis indicated that these genes are enriched for those involved in ADP metabolic processes and cytoplasmic translation. To find out if this was a common trend among highly transcribed genes, we grouped genes by their expression level and determined H3K36me2/me3 levels in each of the groups. Our analysis showed revealed only a slight decrease in H3K36me2/me3 levels in the most highly expressed genes relative to other groups in the *spt6-1004 SET2-H366N* strain (Supplementary Fig. 5g), indicating that the level of transcription was not the determining characteristic among this set of genes. Our results suggest the possibility of a different mechanism for regulation of Set2 activity at these genes.

### The Set2-Spt6 genetic interaction is conserved

As both Set2 and Spt6 are conserved, including the Set2 autoinhibitory domain (Fig. 7a; ^47^), we wanted to test whether the functional interactions between Set2 and Spt6 are conserved. To test this idea, we moved to *S. pombe*, a yeast that is as evolutionarily diverged from *S. cerevisiae* as either is from mammals^51^. We construced an *S. pombe* strain that contains a *set2* mutation similar to the *S. cerevisiae SET2Δ3* mutation (Fig. 7a) and asked whether it could suppress the H3K36 methylation defect caused by an *S. pombe spt6* mutation. For these experiments, we used an *S. pombe spt6-1* mutant which, like *S. cerevisiae spt6-1004*, has a deletion of the sequence encoding the Spt6 HhH domain^24^. This mutant has no detectable H3K36me2 or H3K36me3, while maintaining normal Set2 protein levels^24^. Our results show that the *S. pombe set2Δ3* mutation suppressed the H3K36me2 defect in *spt6-1* although not the H3K36me3 defect (Fig. 7b-e). The lack of suppression of the H3K36me3 defect is likely related to our finding that in a wild-type *spt6^+^* background, the *S. pombe set2Δ3* mutation caused decreased levels H3K36me3 compared to wild-type, suggesting some functional differences for Set2 between the two species^52^. In spite of these differences, our results show that the autoinhibitory domain region of Set2 and its functional interaction with Spt6 is conserved between the two distantly-related yeasts.

**Fig. 7.**
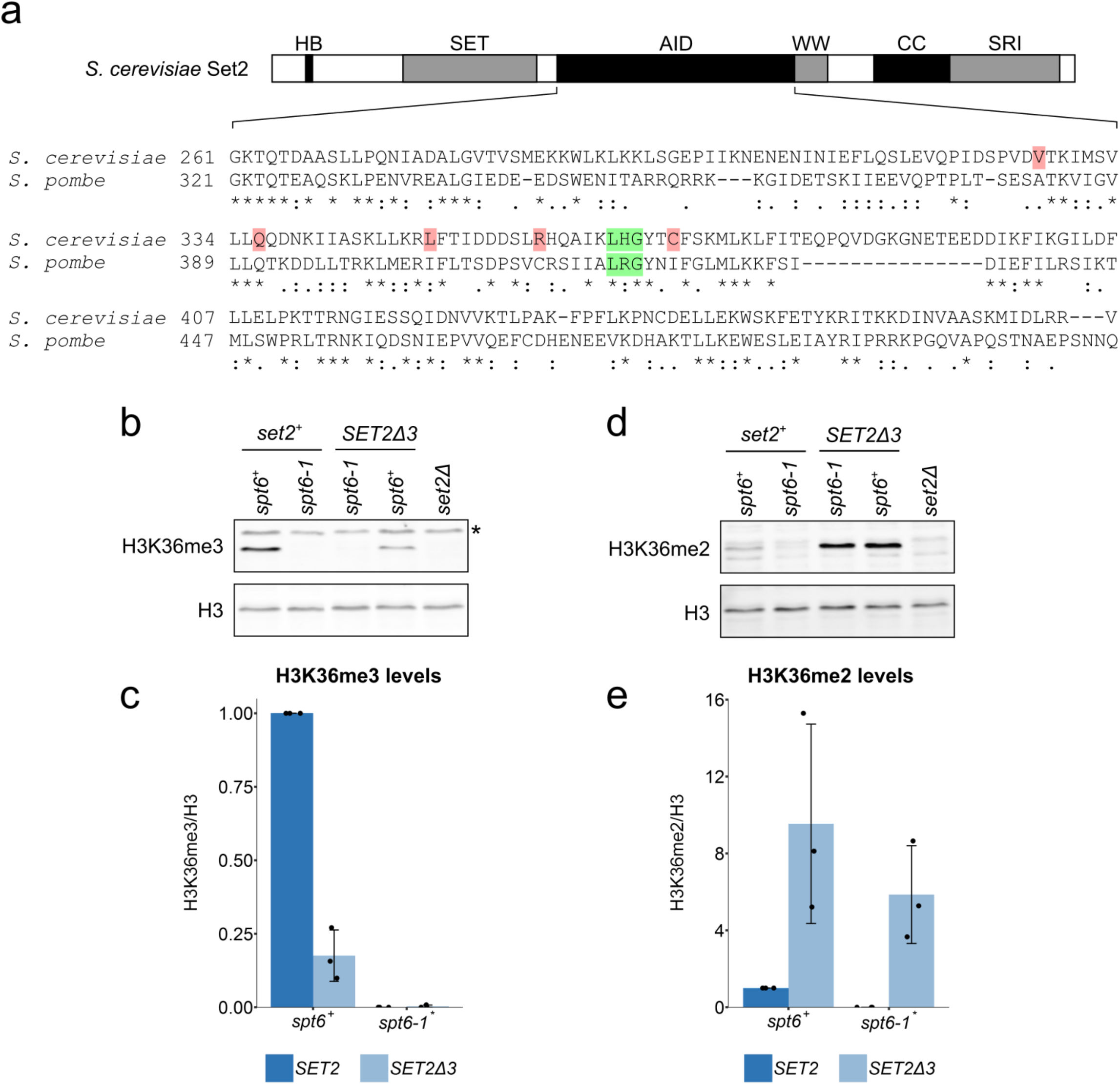
The genetic interaction between Spt6 and Set2 is conserved in *S. pombe*. **a**, Conservation of the amino acid sequence of the central region of Set2 between *S. cerevisiae* (amino acids 261-475) and *S. pombe*. The residues highlighted in green correspond to the three amino acids deleted in the *SET2Δ3* mutation. The residues highlighted in pink denote the location of the other *SET2^sup^* mutations. An asterisk indicates an identical residue, a colon indicates a highly similar amino acid (scoring > 0.5 in the Gonnet PAM 250 matrix), and a period indicates a weakly similar amino acid (scoring <= 0.5 in the Gonnet PAM 250 matrix). **b**, Western blot assaying the effect of the *set2Δ3* mutation on H3K36me3 levels in *spt6-1* cells in *S. pombe*. Histone H3 was used as a loading control. Asterisk denotes a non-specific band. **c**, Quantification of Western blots assaying H3K36me3 levels in the indicated strains. **d**, Western blot assaying the effect of the *set2Δ3* mutation on H3K36me2 levels in *spt6-1* cells in *S. pombe*. Histone H3 was used as a loading control. **e**, Quantification of Western blots assaying H3K36me2 levels in the indicated strains. In both graphs, the black dots represent the individual data points for three experiments and the bars show the mean +/- standard deviation.

## Discussion

In this work we have presented new results about the regulation of Set2 activity in *S. cerevisiae*, providing insights into the requirements for Set2 to function during transcription elongation. By the isolation of *SET2^sup^* mutations that partially suppress the requirement for Spt6, we have shown that a recently identified Set2 autoinhibitory domain^47^ plays critical roles in the regulation of Set2 *in vivo*. Our results also suggest that the Set2 autoinhibitory domain dictates that Set2 will only be active in the presence of Spt6 and the PAF complex. Furthermore, our results also suggest that the Set2 autoinhibitory domain requires that Set2 interacts with both RNAPII and histones via the Set2 SRI and HB domains, respectively, in order to function. We have demonstrated that the dependence of Set2 on Spt6 occurs genome-wide, at a step subsequent to the recruitment of Set2 to chromatin. Together, these results suggest a model in which the Set2 autoinhibitory domain evaluates multiple interactions between trans-acting factors and specific Set2 domains before allowing Set2 to catalyze H3K36me2/me3 *in vivo*. If any of the interactions fail to occur, then the Set2 autoinhibitory domain inhibits Set2 catalytic activity.

A majority of the single amino acid changes (18/20) identified by our *SET2^sup^* mutations fall within a predicted single alpha helix of a proposed Set2 autoinhibitory domain^47^. The tight clustering of our mutations likely reflects the stringency of our mutant selection. The nature of the mutations that we isolated suggests that they disrupt the alpha helix, thereby impairing the autoinhibitory domain, as seven of the 20 mutations encode proline. Furthermore, a deletion of the three codons that contained 13 of the 20 mutations also confers the same phenotype. The previous work that showed that deletion mutations spanning a four-helix region resulted in hyperactive Set2 proteins both *in vitr*o and *in vivo*, although the *in vivo* analysis was limited by the instability of the mutant proteins^47^. The *SET2^sup^* mutations that we have isolated encode stable proteins, which allowed us to discover the critical role of the autoinhibitory domain *in vivo*.

Our results raise the question of the mechanism by which Spt6 is required for Set2 activity. At 30°C, the temperature at which our experiments were performed, the Spt6-1004 mutant protein is present at normal levels (N.I. Reim and F. Winston, unpublished results), yet there is no detectable H3K36me2/me3. The simplest possibility is that a direct interaction between the Spt6 HhH domain, the region missing in the Spt6-1004 mutant protein, and the Set2 autoinhibitory domain is required for Set2 activity. In support of this idea, the *spt6-1004* mutation causes the most severe defects in H3K36me2/me3 of all *spt6* alleles tested^18,50^. However, there is no evidence for a direct Spt6-Set2 interaction, either by high-resolution analysis of Set2-interacting proteins^53^ or by two-hybrid analysis (our unpublished results). While these negative results do not rule out a direct Set2-Spt6 interaction, it also seems plausible that in the *spt6-1004* mutant there is an altered chromatin configuration that impairs the interaction of Set2 with a nucleosomal surface post-recruitment, such as that previously identified to be required for Set2 activity^42,43^.

Although the *SET2^sup^* mutations are able to suppress the *spt6-1004* H3K36 methylation defect for most genes, there are a small number of genes at which H3K36 methylation is not rescued. This finding suggests that the mechanisms that regulate Set2 activity *in vivo* may not be uniform across the genome. The genes that behaved differently are highly transcribed and have a lower level of histone H3 compared to most genes; however, neither of those characteristics is sufficient to explain their lack of response to the *SET2^sup^* mutations. At these genes, there may be additional or distinct requirements for Set2 to function. Alternatively, these genes may recruit a high level of H3K36 demethylases or, as H3K36 can also be acetylated^54,55^, these genes may be more subject to competition between these mutually exclusive modifications than at most other genes.

Autoinhibition is a common mode of regulation among histone methyltransferases in both yeast and mammals. For example, the *S. cerevisiae* H3K4 methyltransferase Set1 also contains an autoinhibitory domain. Point mutations that alter this domain in Set1, similar to mutations in the Set2 autoinhibitory domain, make Set1 hyperactive and independent of transcription elongation factors that are normally required for its activity^56^. In addition, mammalian histone methyltransferases have been shown to contain autoinhibitory domains, including Nsd1^57^, PRDM9^58^, and Smyd3^59^. In these three cases, structural studies of the proteins have suggested likely mechanisms that are distinct from each other. There are at least two reasons that a methyltransferase such as Set2 would have such tight regulation of its activity. First, the requirement for interactions with both nucleosomes and elongating RNAPII, as well as the activities of factors such as Spt6 and Paf1, ensures that H3K36me2/me3 will only occur at the correct location and at the correct time – on chromatin when it is being actively transcribed. Second, this regulation provides the opportunity to regulate H3K36me2/me3 in different conditions. For example, recent studies have provided evidence that H3K36 methylation is important for nutrient stress response^31,60^, carbon source shifts^61^, DNA damage responses^62–65^, splicing^66–69^, and aging^70,71^. Therefore, the Set2 autoinhibitory domain may serve as a target for additional regulators under particular growth conditions.

## Methods

### Yeast strains and media

All *S. cerevisiae* and *S. pombe* strains used in this study were constructed by standard methods and are listed in Table S2. The *S. pombe set2Δ3* mutation was made based on alignment of the *S. cerevisiae* and *S. pombe* Set2 amino acid sequence using the Uniprot ‘Align’ tool (https://www.uniprot.org/help/sequence-alignments.) All *S. cerevisiae* liquid cultures were grown in YPD (1% yeast extract, 2% peptone and 2% glucose) at 30°C unless mentioned otherwise. All *S. pombe* liquid cultures were grown in YES (0.5% yeast extract, 3% glucose, 225 mg/l each of adenine, histidine, leucine, uracil, and lysine) at 32°C. All strains were constructed using transformations and/or crosses. For the genetic selection, the two reporter genes were constructed individually and then crossed to each other. The *FLO8-URA3* reporter was constructed by inserting the *URA3* gene at the 3’ end of the *FLO8* gene, replacing base pairs +1727 - +2505 (+1 = ATG)^48^. The *STE11-CAN1* reporter was constructed by inserting the *CAN1* gene at the 3’ of the *STE11* gene, replacing base pairs +1871 - +2154 (+1 = ATG)^48^. In the same strain, the coding sequence of the endogenous *CAN1* gene was deleted using HygMX cassette, which was amplified from the *pFA6a-hphMX6* plasmid^72^. To make the strain containing the *STE11-CAN1* reporter amenable to crosses, the *STE11-TAP-HIS3MX* cassette was amplified from a strain derived from the yeast TAP-tagged collection^73^ and inserted at the *HO* locus replacing base pairs −1400 to +1761 (+1 = ATG). For spot tests to check for reporter expression, cells were spotted on media containing 1mg/ml 5-FOA and/or 150 μg/ml canavanine unless mentioned otherwise. For verification of mutants obtained from the selection, point mutations were made in the *SET2* gene by two step gene replacement^74^ in a strain containing only the *STE11-CAN1* reporter. The same strategy was used to make the different deletions within the region of the *SET2* gene that codes for the autoinhibitory domain. For Spt6 depletion experiments, cells were grown to OD_600_ ≈ 0.6 (~2×10^7^ cells). Cells were diluted in YPD to OD_600_ = 0.3. A fraction of the diluted culture was collected and treated as the 0 minute time point. Indole acetic acid (IAA) dissolved in DMSO was then added to the medium at a final concentration of 25 μM and cultures were grown at 30°C. Cells were collected at 90, 120 and 150 minutes to prepare whole cell extracts for Western blotting.

### Isolation and analysis of mutants that suppress intragenic transcription

The genetic selection for isolating mutants that suppress intragenic transcription in *spt6-1004* was done in two rounds. In the first round, 25 independent cultures each of FY3129 and FY3130 were grown to saturation overnight in YPD. Cells were washed twice with water and 2-4 × 10^7^ cells from each independent culture were spread on two SC-Arg plates containing 0.25 mg/ml 5-FOA and 150 μg/ml canavanine, one of which was UV irradiated. All plates were incubated at 34°C, which reduced background growth, and colonies that grew between days 3-7 were picked for further analysis. The second round of selection was identical to the first round, except that cells were plated on SC-Arg plates containing 0.5 mg/ml 5-FOA and 150 μg/ml canavanine. To test for dominance, the suppressor strains were first crossed with the parent *spt6-1004* strain carrying both the reporters, diploids were selected by complementation, and the purified diploids were tested for growth on medium containing 5-FOA and canavanine. To test for linkage, suppressors were crossed to each other, sporulated, and tetrads were dissected and analyzed by standard conditions. At least 10 tetrads were analyzed per cross.

### Identification of suppressor mutations by whole-genome sequencing

Eight of the twenty independent dominant mutants identified were subjected to further genetic analysis and found to contain mutations in a single gene responsible for the suppression phenotype. Whole genome sequencing libraries were prepared for the eight mutants and two parents (RGC95 and RGC98) as described previously^75^. Sonication of genomic DNA was done using Covaris S2 (3 cycles of 50"; Duty cycle: 10%; Intensity: 4; Cycles/burst: 200) to obtain fragments between 100-500 bp. GeneRead DNA library prep kit (QIAGEN) was used for end repair, A tailing and adapter ligation. The DNA samples were purified twice using 0.7x volume SPRI beads. PCR cycles for final amplification were selected based on trial amplification runs. Following the final PCR, DNA was purified twice using 0.7x volume SPRI beads and submitted for next generation sequencing. Sequencing was done on an Illumina Hi-Seq platform. Reads from the FASTQ file were aligned using Bowtie2^76^ (default parameters) and variants between all ten libraries and the reference genome were called using the *samtools mpileup* command^77^. The resulting VCF file was then used to identify variants that were present only in the mutants and not in the parents using a custom R script (https://github.com/winston-lab). All variants that mapped within the coding sequence of a gene were identified. The gene that had variants in all mutants was found to be *SET2*.

### Spot tests

Yeast cultures were grown overnight from single colonies. Cells were pelleted and washed once with water. All cultures were normalized by their OD_600_ values and six ten-fold serial dilutions of each culture were made in a 96-well plate. The cultures were spotted on different media and incubated at the appropriate temperatures. All plates containing FOA and/or canavanine were incubated at 34°C to ensure higher stringency for assaying intragenic transcription. The control complete plates for these experiments were also incubated at 34°C. The Spt- phenotype was assayed at 30°C. Temperature sensitivity was assayed at 37°C. For mutant backgrounds with weak intragenic transcription (as in Fig. 4), expression of the *FLO8-URA3* reporter was tested on SC-Ura medium.

### Western blotting and antibodies

*S. cerevisiae* cells were grown to OD_600_ ≈ 0.6 (~2×10^7^ cells). Culture volumes were normalized by their OD_600_ such that an OD equivalent (OD_600_ * volume of the culture) of 6 was harvested. The cell pellets were washed once with water and suspended in 300 μl water. Then, 300 μl of 0.6M sodium hydroxide was then added to the cells and the suspension was incubated at room temperature for 10 minutes. The cells were then pelleted and resuspended in 80 μl Modified SDS buffer (60mM Tris–HCl, pH 6.8, 4% β-mercaptethanol, 4% SDS, 0.01% bromophenol blue, and 20% glycerol)^78^. Eight microliters of the extracts were loaded on an SDS-PAGE gel for western blotting. *S. pombe* cells were grown to OD_600_ ≈ 0.6 (~10^7^ cells/ml). 10 ml of cells were pelleted and resuspended in 200 μl of 20% tri-chloroacteric acid (TCA). Next, 200 μl of glass beads were added to the tube and the cells were lysed by bead beating for 2 minutes at 4°C. The bottom of the tube was punctured and the flow through collected by centrifugation. The beads were washed twice with 200 μl of 5% TCA and the flow through was collected. The pooled flow through fractions were spun at 3000 rpm for 10 minutes at room temperature and the resulting pellet was resuspended in 150 μl (normalized by OD_600_; culture of OD_600_ = 0.8 was suspended in 150 μl) of 2x Laemmli buffer (125 mM Tris-HCl pH 6.8, 4% SDS, 10% β-mercaptoethanol, 0.01% bromophenol blue, 40% glycerol). An equal volume of 1M Tris base (pH not adjusted) was added to neutralize the TCA. The samples were incubated for 5 minutes at 95°C and spun down at 10,000 rpm for 30 seconds. Then, 10 μl of the supernatant was loaded on an SDS-PAGE gel for western blotting. Primary antibodies used for Western blotting were: anti-Set2 (1:8000, generously provided by Brian Strahl), anti-H3K36me3 (1:2000, Abcam, ab9050), anti-H3K36me2 (1:2500, Abcam, ab9049) or (1:1000, Upstate #07-274), anti-HA (1:5000, Abcam, ab9110), anti-Flag (1:5000, Sigma, F3165), anti-Spt6 (generously provided by Tim Formosa), anti-H3 (1:2500, Abcam, ab1791), anti-Pgk1 (1:10000, Life Technologies 459250), and anti-Act1 (1:10000, Abcam, ab8224). Secondary antibodies used were: goat anti-rabbit IgG (1:10000, Licor IRDye 680RD) and goat anti-mouse IgG (1:20000, Licor, 800CW). Quantification of Western blots was done using Licor ImageStudio software.

### Northern blotting

RNA extraction from *S. cerevisiae* was done using hot acid phenol extraction as described previously^79^. Northern blotting was done as described previously^79^ with many modifications. Fifteen μg of RNA was loaded per sample. The composition of the final RNA loading dye was 6% formaldehyde, 1x MOPS, 2.5% Ficoll, 10mM Tris-HCl, pH 7.5, 10mM EDTA, 7 μg/ml ethidium bromide, 0.025% Bromophenol blue and 0.025% Orange G. Following the addition of RNA loading dye, the RNA sample was heated at 65°C for 5 minutes and then transferred to ice before loading on the gel. Transfer of the RNA from gel to the membrane was done using upward capillary transfer in 1x SSC solution. Pre-hybridization of the membrane was done for 3-5 hours in pre-hybridization solution (50% deionized formamide, 10% dextran sulphate, 1M NaCl, 0.05M Tris-HCl pH 7.5, 0.1% SDS, 0.1% sodium pyrophosphate, 10x Denhardts reagent, 500 μg/ml denatured salmon sperm DNA) at 42°C. Following hybridization, six washes were done - 2 washes with 2x SSC solution at room temperature for 15 minutes each, 2 washes with 2x SSC, 0.5% SDS at 65°C for 30 minutes each, and 2 washes with 0.1x SSC at room temperature for 30 minutes each. Probes were made with the PCR primers listed in Table S3.

### ChIP and ChIP-Seq

For *S. cerevisiae* cultures, 140 ml of cells were grown to OD_600_ ≈ 0.6 (~2×10^7^ cells) in YPD. Cultures were cross-linked by the addition of formaldehyde to a final concentration of 1% followed by incubation with shaking at room temperature for 20 minutes. Glycine was added to a final concentration of 125 mM and the incubation was continued for 10 minutes. The cells were pelleted and washed twice with cold 1x TBS (100mM Tris, 150 mM NaCl, pH 7.5) and once with cold water. The cell pellets were then suspended in 800 μl cold LB140 buffer (50 mM HEPESKOH, pH 7.5, 140 mM NaCl, 1mM EDTA, 1% Triton X-100, 0.1% sodium deoxycholate, 0.1% SDS, 1x cOMPLETE Protease Inhibitor tablet (Roche)). One ml of glass beads was added and the cells were lysed by bead beating for 8 minutes at 4°C with incubation on ice for 3 minutes after every one minute. The lysate was collected and centrifuged at 12,500 rpm for 5 minutes and the resulting pellet was washed once with 800 μl cold LB140 buffer. The pellet was resuspended in 580 μl cold LB140 buffer and sonicated in a QSonica Q800R machine for 20 minutes (30 seconds on, 30 seconds off, 70% amplitude). The sonicated samples were centrifuged at 12,500 rpm for 30 minutes and the resulting supernatant was taken for the immunoprecipitation step. For the *S. pombe* spike-in strain, 120 ml of cells were grown to OD_600_ ≈ 0.6 in YES and processed similar to the *S. cerevisiae* culture except for the following steps: Bead beating for cell lysis was done for 11 minutes. Sonication was done for 15 minutes. The protein concentrations in chromatin were measured by Bradford assay^80^. 300-500 μg of *S. cerevisiae* chromatin was mixed with 33-55 μg (10%) of *S. pombe* chromatin and the volume was brought up to 800 μl with WB140 buffer (50 mM HEPES-KOH pH 7.5, 140 mM NaCl, 1mM EDTA, 1% Triton X-100, 0.1% sodium deoxycholate). This was used as the input for the immunoprecipitation reaction. Antibody (amounts mentioned below) was added to the input and the samples incubated overnight at 4°C with end-over-end rotation. Fifty μl of Protein G sepharose beads (GE healthcare) pre-washed twice in WB140 was added to the IPs and samples were incubated for 4 hours at 4°C with end-over-end rotation. The beads were washed twice with WB140, twice with WB500 (50 mM HEPES-KOH, pH 7.5, 500 mM NaCl, 1mM EDTA, 1% Triton X-100, 0.1% sodium deoxycholate), twice with WBLiCl (10 mM Tris pH 7.5, 250 mM LiCl, 1mM EDTA, 0.5% NP-40, 0.5% sodium deoxycholate) for two minutes each and once with TE (10 mM Tris pH 7.4, 1 mM EDTA) for five minutes. The immunoprecipitated material was eluted twice with 100 μl TES (50 mM Tris pH 7.4, 1 mM EDTA, 1% SDS) at 65°C for 30 minutes. The eluates were incubated at 65°C overnight to reverse the crosslinking. Two hundred μl of TE was added to the eluates followed by RNase A/T1 to a final concentration of 0.02 μg/μl. The samples were incubated at 37°C for 2 hours. Proteinase K was added to a final concentration of 0.4 mg/ml and samples were incubated at 42°C for 2 hours. DNA was purified using Zymo DCC (for ChIP-Seq) or EZNA Cycle Pure kit spin columns (for ChIP-qPCR). The purified DNA was used for qPCR or library preparation for next generation sequencing. Primers used for qPCR are listed in Supplementary Table 3. The library preparation steps from this stage were similar to those used for preparation of DNA libraries for whole genome sequencing^75^. Next generation sequencing was done on an Illumina NextSeq platform. Five μl of anti-HA (Abcam, ab9110) per 500 μg of chromatin, 10 μl (5 μl for ChIP-qPCR) of anti-Rpb1 (Covance, 8WG16) per 500 μg of chromatin, 4 μl of anti-H3 (Abcam, ab1791) per 300 μg of chromatin, 4 μl of anti-H3K36me2 (Abcam, ab9049) per 300 μg of chromatin, 4 μl of anti-H3K36me3 (Abcam, ab9050) per 300 μg of chromatin, and 50 μl of anti-FLAG M2 affinity gel (Sigma, #A2220) per 500 μg of chromatin were used for ChIP. HA tagged Set2 was used for all ChIP-seq and follow-up ChIP-qPCR experiments. FLAG tagged Set2 was used for ChIP-qPCR of set2 mutants lacking the SRI and HB domains.

**ChIP-seq computational analysis**

A custom ChIP-seq pipeline was generated using the Snakemake workflow manager^81^. ChIP-seq data was aligned to a combined *S. cerevisiae* + *S. pombe* genome using Bowtie2^76^. 72% - 93% of the reads mapped exactly once to the combined genome. The number of mapped reads for *S. cerevisiae* IPs (excluding inputs) varied from 1 million - 10 million reads. The number of mapped reads for *S. pombe* immunoprecipitation (excluding inputs) varied from 350,000 - 6 million reads. Cross correlation was done using SPP package^82^ to estimate fragment sizes for the different libraries. The coverage files were produced using the igvtools count function^83^, extending reads by (length of fragment sizes - average read length) for each library. Spike in normalization was done as described previously^84^, correcting for variations in the input samples. Correlation plots, heatmaps and metagene plots were produced using custom R scripts that are available upon request. To identify genes that did not show rescue of H3K36me2 in the suppressor strain, coverage within 20 bp windows tiling the entire genome was generated for each library. IP libraries were divided by their respective control libraries after the addition of a pseudocount of one. The mean coverage over every gene in each library was determined using the bedtools map command^85^. The ratio of the mean coverage for every gene in one sample over the other was calculated.

### Data availability

Genomic datasets are deposited in the Gene Expression Omnibus with accession number GSE116646. Other primary data are available from the corresponding author upon reasonable request.

## Acknowledgments

We are grateful to Steve Buratowski, Steve Doris, and Catherine Weiner for helpful comments on the manuscript, to Tim Formosa and Brian Strahl for providing antibodies, to Vanessa Cheung for constructing the *FLO8-URA3* reporter, and to Natalia Reim for providing the Spt6 degron strain. This work was supported by NIH grant GM032967 to F.W.

## Author contributions

R.G. and F.W. conceived of the experiments. R.G. performed the experiments and the data analysis. R.G. and F.W. wrote the manuscript.

---

**Additional information** Supplementary information is available for this paper at

**Supplementary Fig. 1.**
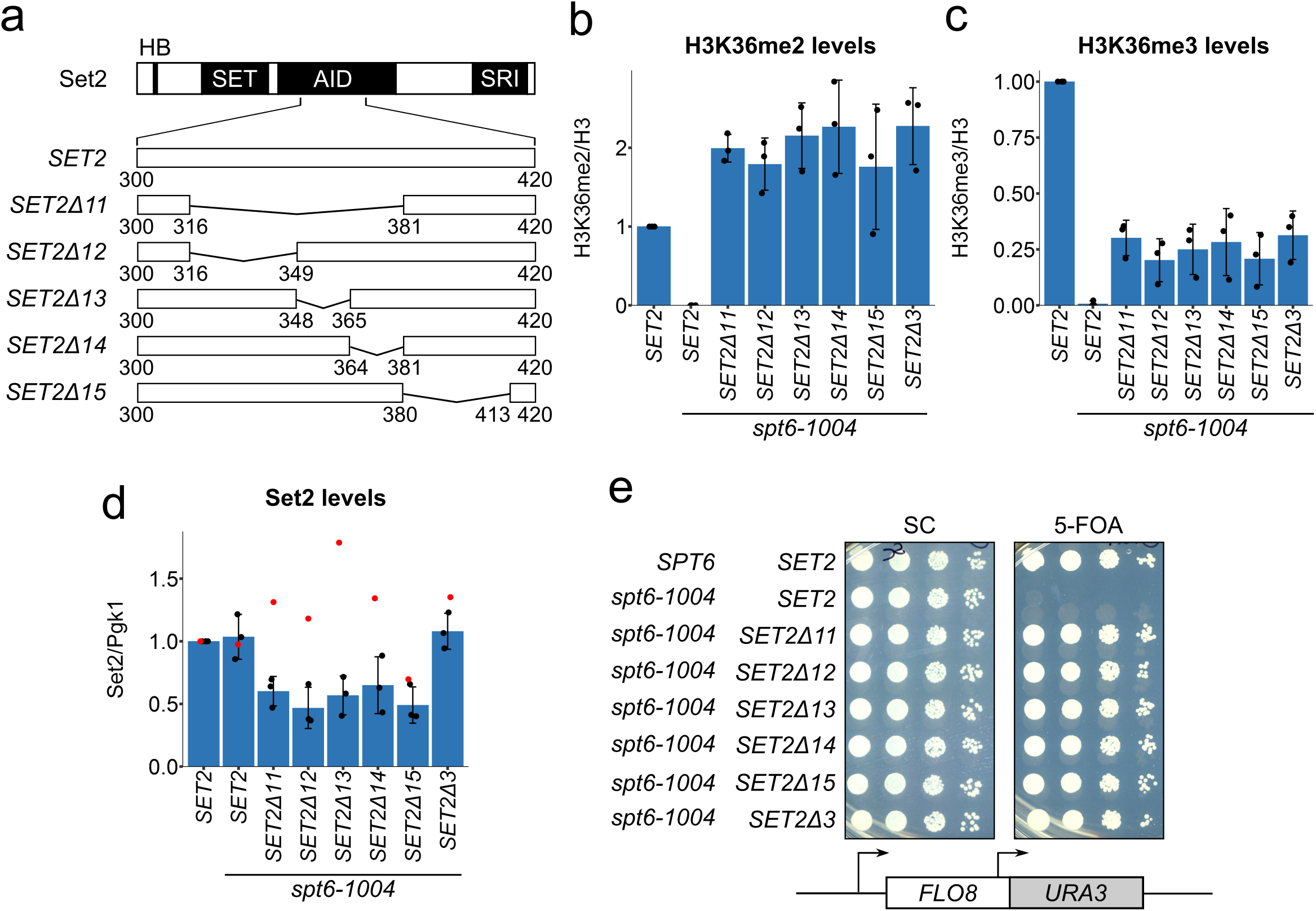
Deletions in the region encoding the Set2 autoinhibitory domain partially rescue H3K36 méthylation and intrageni transcription in *spt6-1004*. **a**, The schematic depicts some of the domains in Set2 and the five mutants that have deletions in the region encoding the autoinhibitory domain. **b,c,d**, Quantification of western blots assaying H3K36me2 (b), H3K36me3 (c), and Set2 (d) levels in *spt6-1004* strains with the indicated *SET2* mutations. The black dots represent the individual data points for three experiments and the bars show the mean +/- standard deviation. The red dots in (d) represent outlier values from one experiment which were not included in the calculation of the mean or error bars, **e**, Spot tests of cells grown at 34°C assaying the effect of *SET2* mutations on expression of the *FL08-URA3* reporter in an *spt6-1004* background.

**Supplementary Fig. 2.**
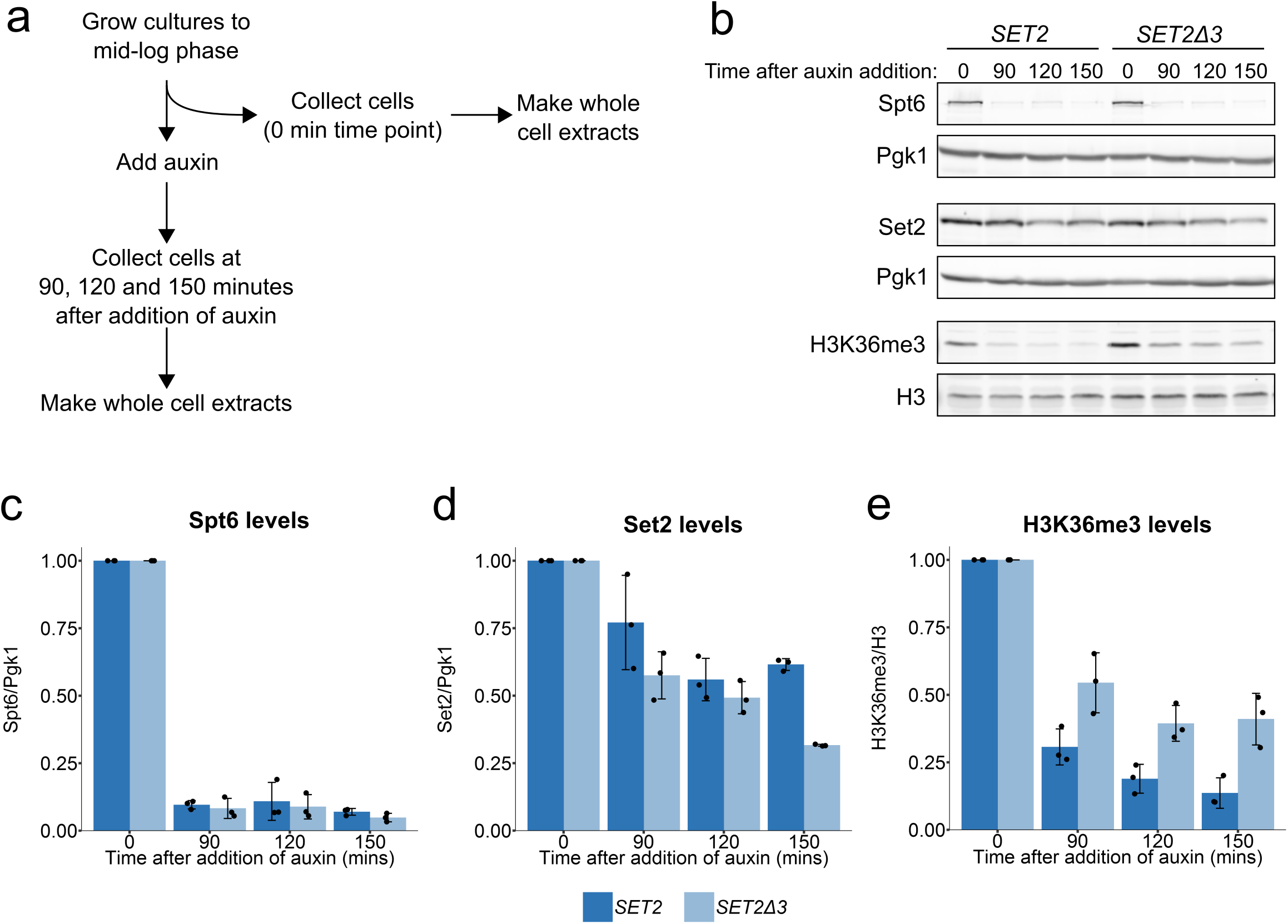
*SET2Δ3* cells show reduced loss of H3K36me3 upon Spt6 depletion. **a**, A schematic illustrating the experimental plan for depletion of Spt6 in wild-type and *SET2Δ3* strains containing a derivative of *SPT6* fused to sequences encoding an auxin-inducible degron. Addition of the auxin IAA promotes degradation of Spt6. **b**, Western blots assaying Spt6, Set2, and H3K36me3 levels in wild-type and *SET2Δ3* strains upon Spt6 depletion. Pgk1 and histone H3 were used as loading controls. **c,d,e**, Quantification of western blots assaying Spt6 (c), Set2 (d) and H3K36me3 (e) levels upon Spt6 depletion, with and without the *SET2Δ3* mutation, normalized to their respective loading controls. The black dots represent the individual data points for three experiments and the bars show the mean +/- standard deviation. Proteins levels for each strain have been normalized to the 0 minute time point.

**Supplementary Fig. 3.**
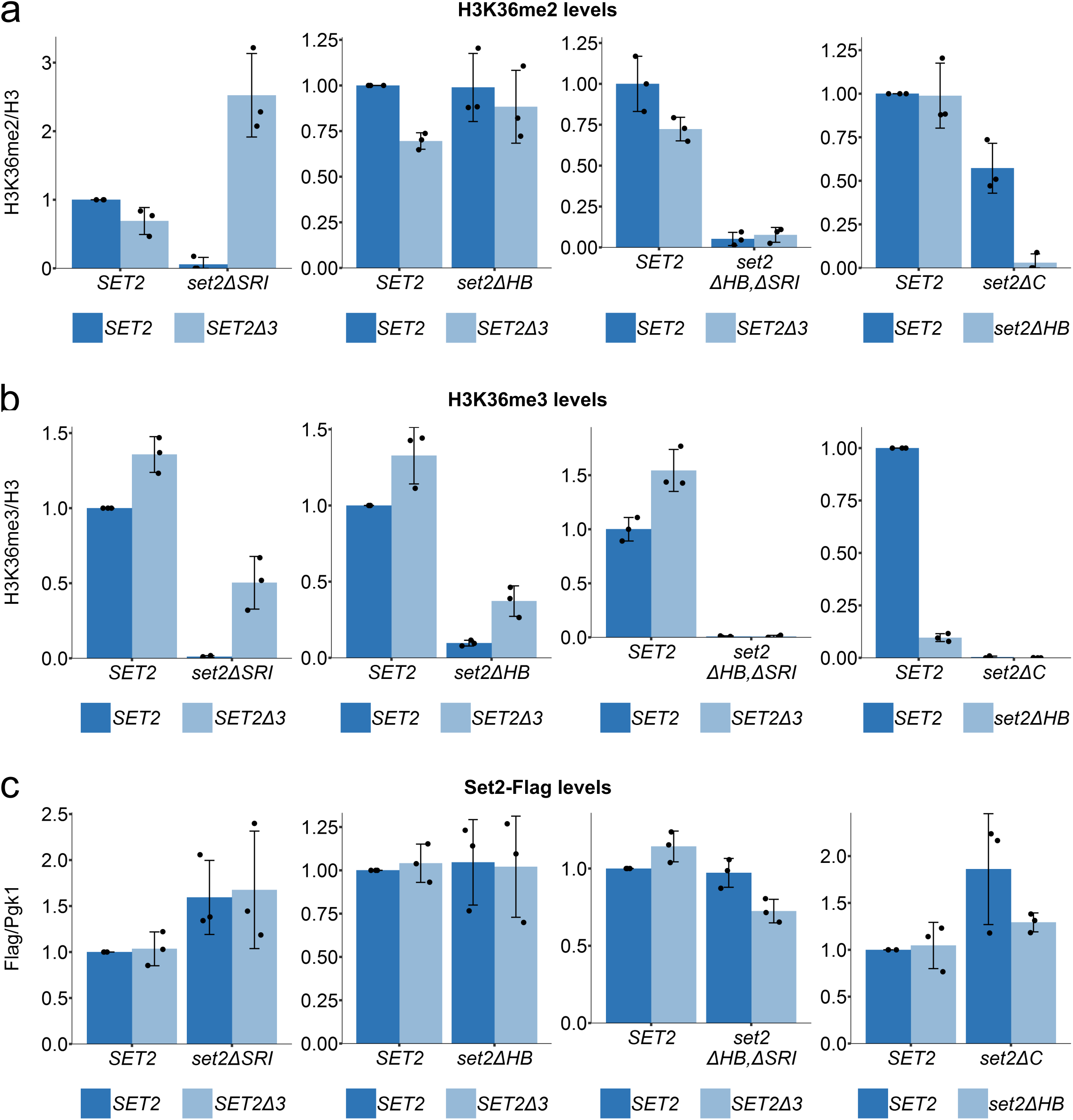
*SET2Δ3* rescues H3K36 méthylation and intragenic transcription upon deletion of the SRI or HB domains. **a,b,c** Quantification of western blots assaying H3K36me2 (a), H3K36me3 (b) and FLAG-tagged Set2 levels (c) in strains with the indicated *set2* mutations. The black dots represent the individual data points for three experiments and the bars show the mean +/- standard deviation. The plotted values for *SET2, SET2Δ3* and *set2ΔHB* are the same between some of the bargraphs, but have been plotted again for ease of representation.

**Supplementary Fig. 4.**
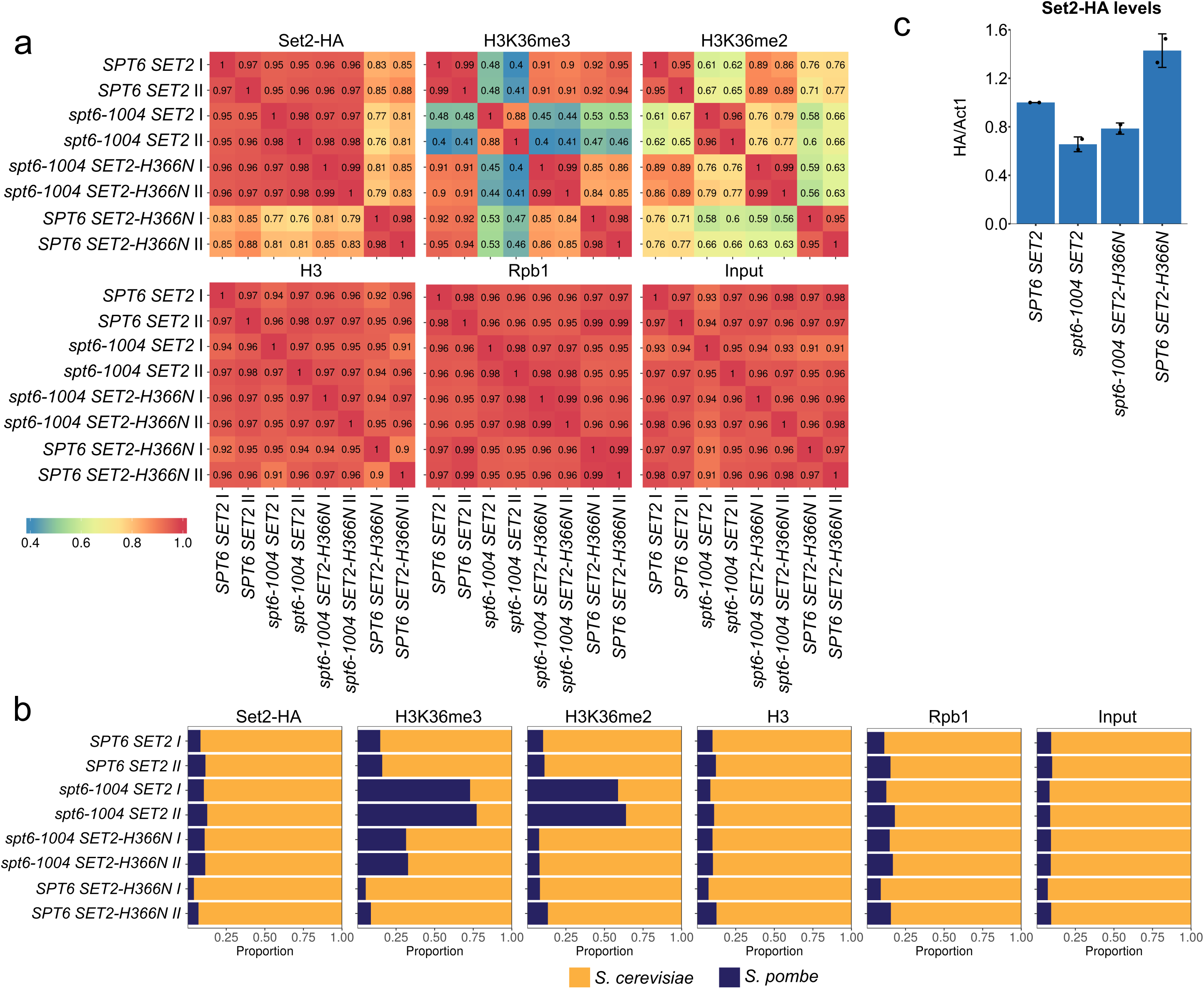
Quality analyses of ChlP-Seq data. **a**, Correlation heatmaps for all 48 samples that were processed for ChlP-Seq. The numbers represent the Pearson correlation coefficient of coverage over 20 bp bins tiling the entire genome, **b**, Bargraphs showing the proportion of reads in each library mapped to the *S. cerevisiae* or *S. pombe* (spike-in control) genome, **c**, Quantification of western blots assaying HA tagged Set2 protein levels prepared from the same cultures that were processed for ChlP-Seq. The black dots represent the individual data points for two experiments and the bars show the mean +/- standard deviation.

**Supplementary Fig. 5.**
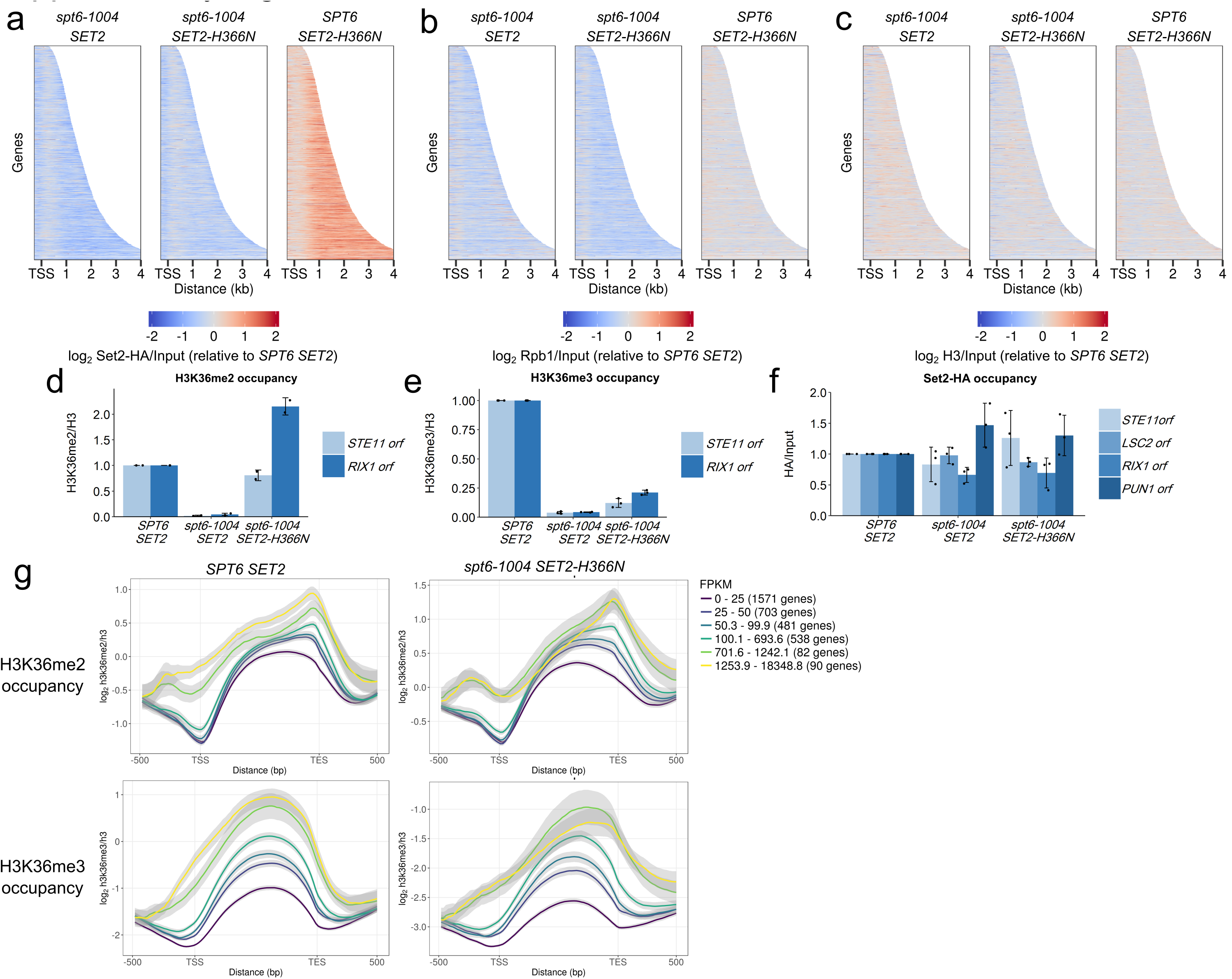
Control analyses for ChlP-Seq data. **a,b,c**, Heatmaps depicting Set2-HA/lnput (a), Rpb1/lnput (b) and H3/lnput (c) levels for all non-overlapping genes that code for protein (n = 3522). All values that had a log fold change of below −2 or above 2 have been set to −2 or 2 respectively. **d,e,f**, ChlP-qPCR analysis quantifying H3K36me2 (d), H3K36me3 (e) and Set2-HA(f) levels over the indicated genes. ChlP-qPCR data has been spike-in normalized to the *S. pombe ACT1* gene, **g**, Metagene plots showing average H3K36me2/H3 and H3K36me3/H3 levels for groups of genes that are binned by their level of expression (obtained from RNA-seq data; N. I. Reim and F. Winston

**Table S1.** *SET2* mutations.

| **Mutation alias** | **Position of Mutation (1=ATG)** | **SET2 mutation** | **Set2 amino acid change** |
| --- | --- | --- | --- |
| LF41α | 979 - 980 | GT -> AA | V327K |
| LF12a | 1006 | C -> G | Q336E |
| LF38α | 1049 | T -> C | L350P |
| LF3α | 1049 | T -> C | L350P |
| HF3α | 1049 | T -> C | L350P |
| LF6a | 1076 | G -> C | R359P |
| S9 | 1094 | T -> A | L365Q |
| LF9α | 1094 | T -> A | L365Q |
| LF39α | 1094 | T -> C | L365P |
| LF43a | 1094 - 1096 | TAC -> CCT | L365P, H366Y |
| S1 | 1096 | C -> A | H366N |
| S7 | 1096 | C -> T | H366Y |
| LF4α | 1096 | C -> T | H366Y |
| HF9α | 1096 | C -> T | H366Y |
| LF7α | 1097 | A -> C | H366P |
| HF6a | 1098 | C -> A | H366Q |
| LF5a | 1099 | G -> A | G367S |
| HF5α | 1099 | G -> T | G367C |
| HF11α | 1100 | G -> A | G367D |
| S3 | 1109 | G -> T | C370F |

**Table S2.**
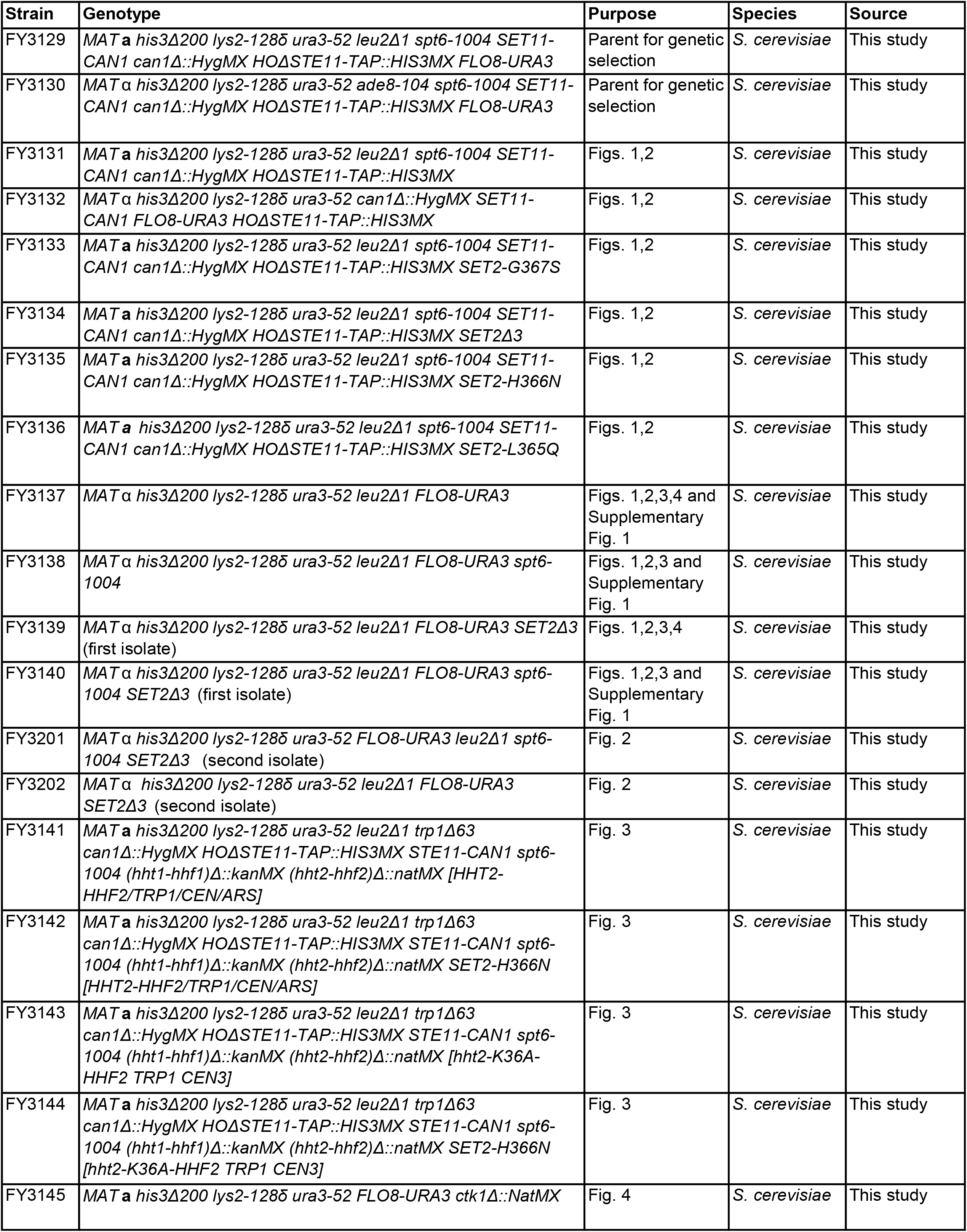
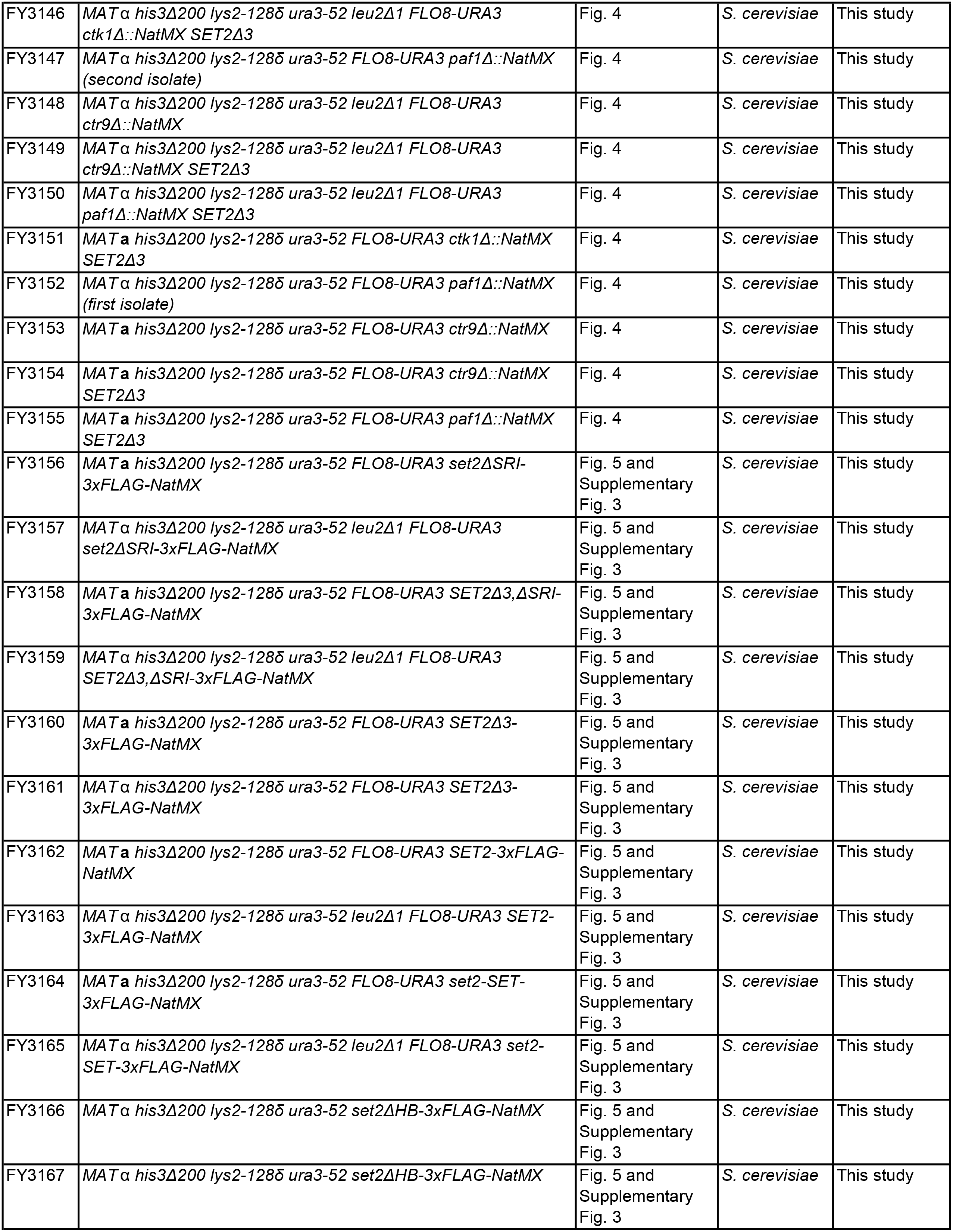
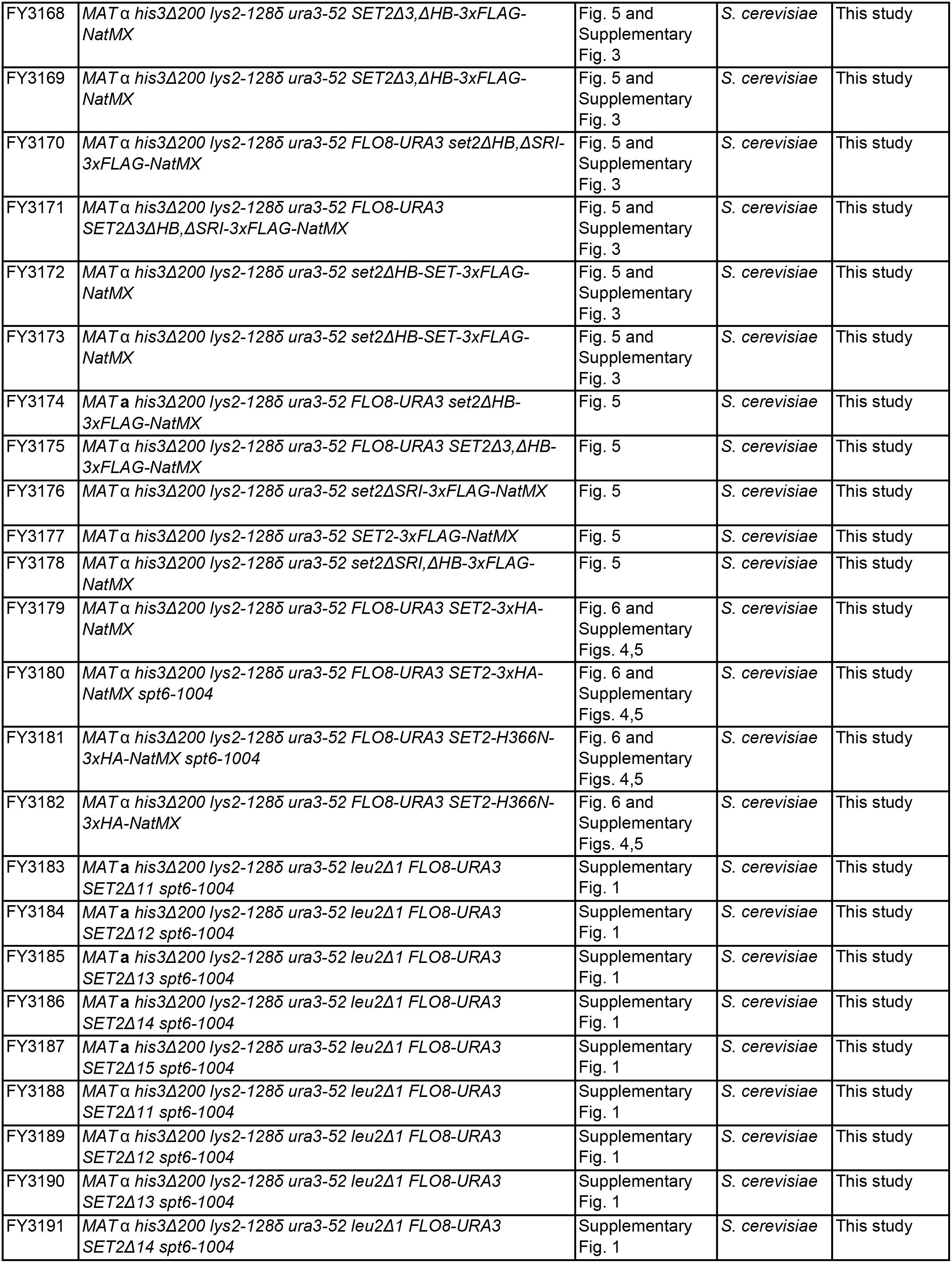
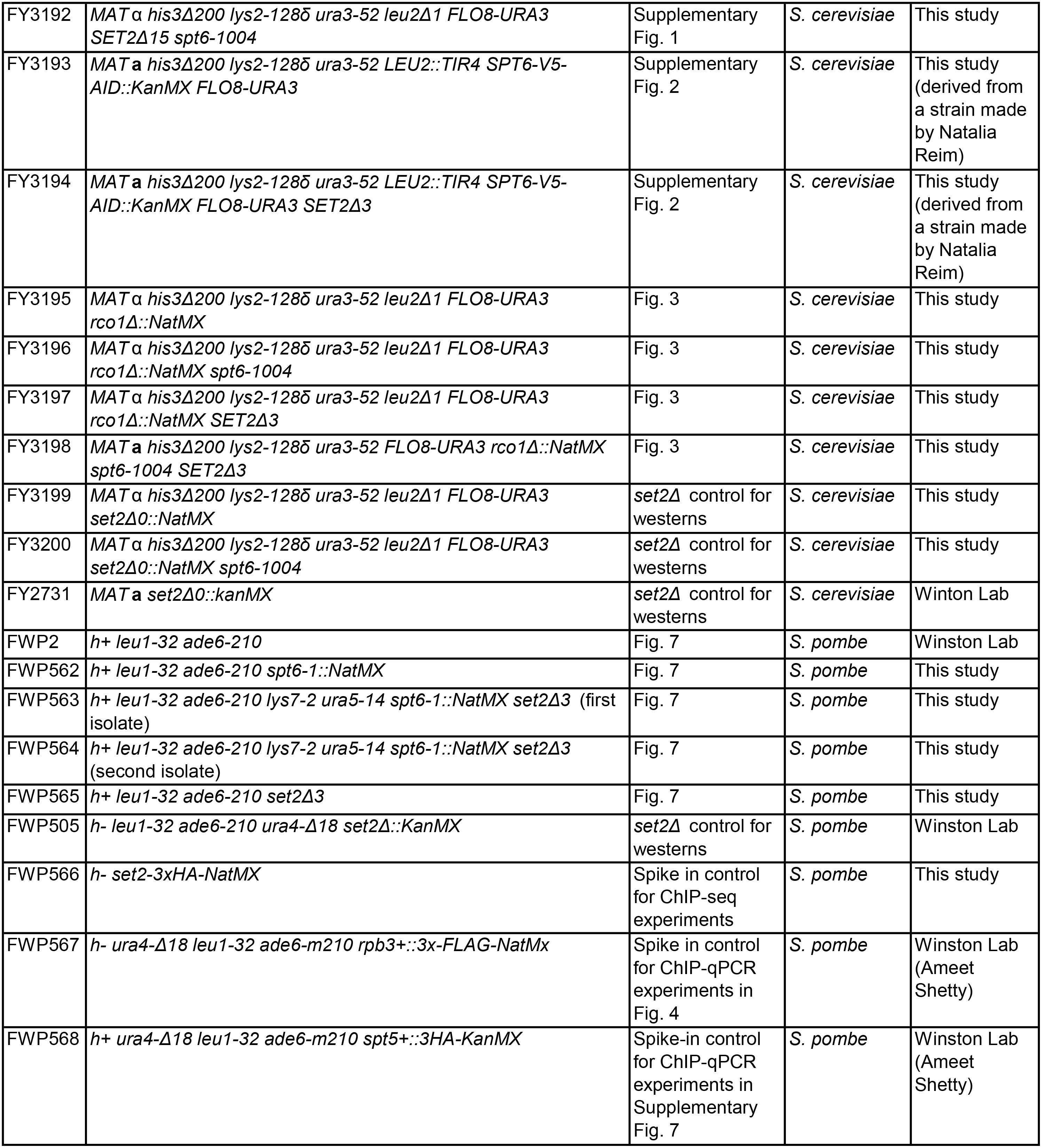

**Table S3.**
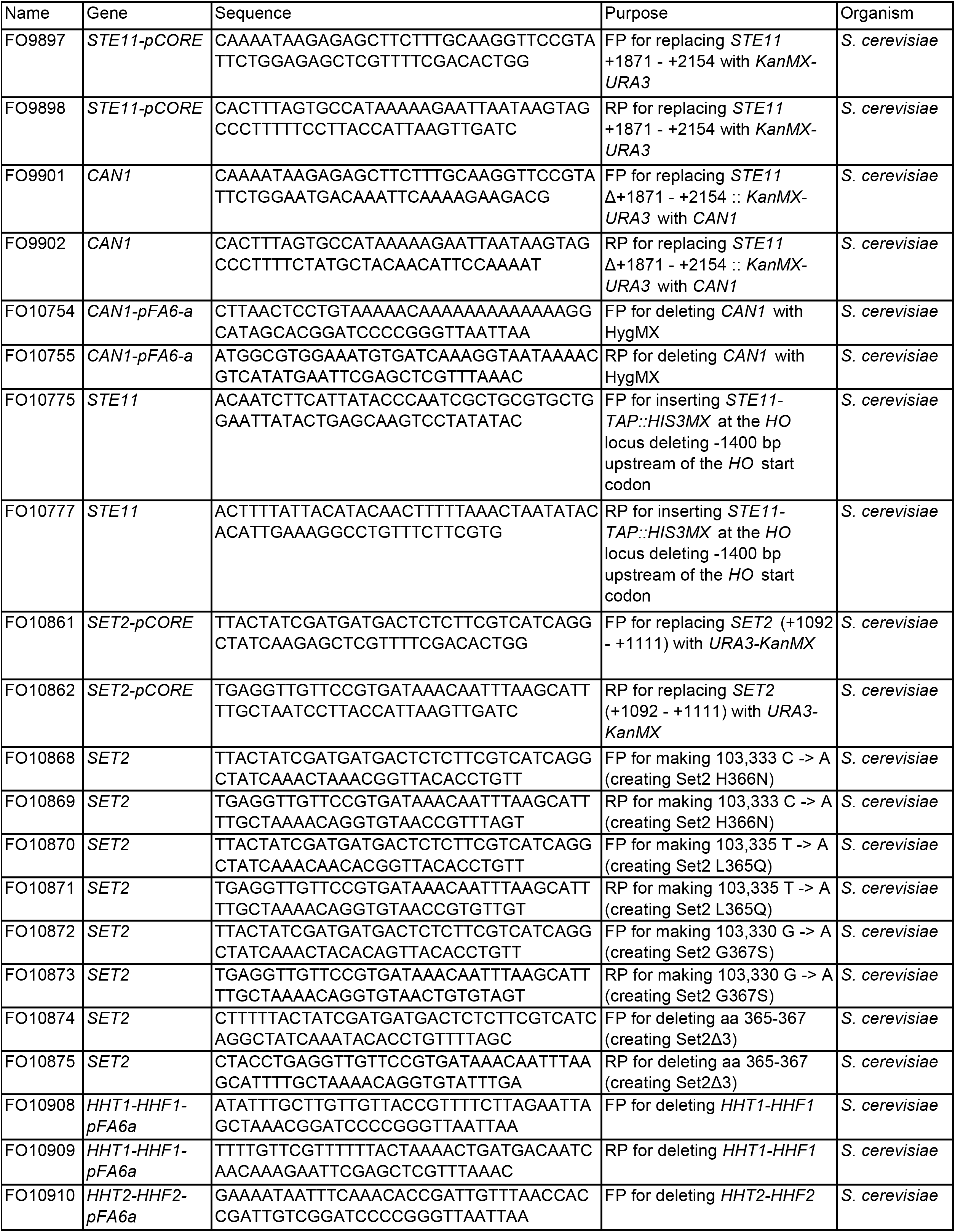
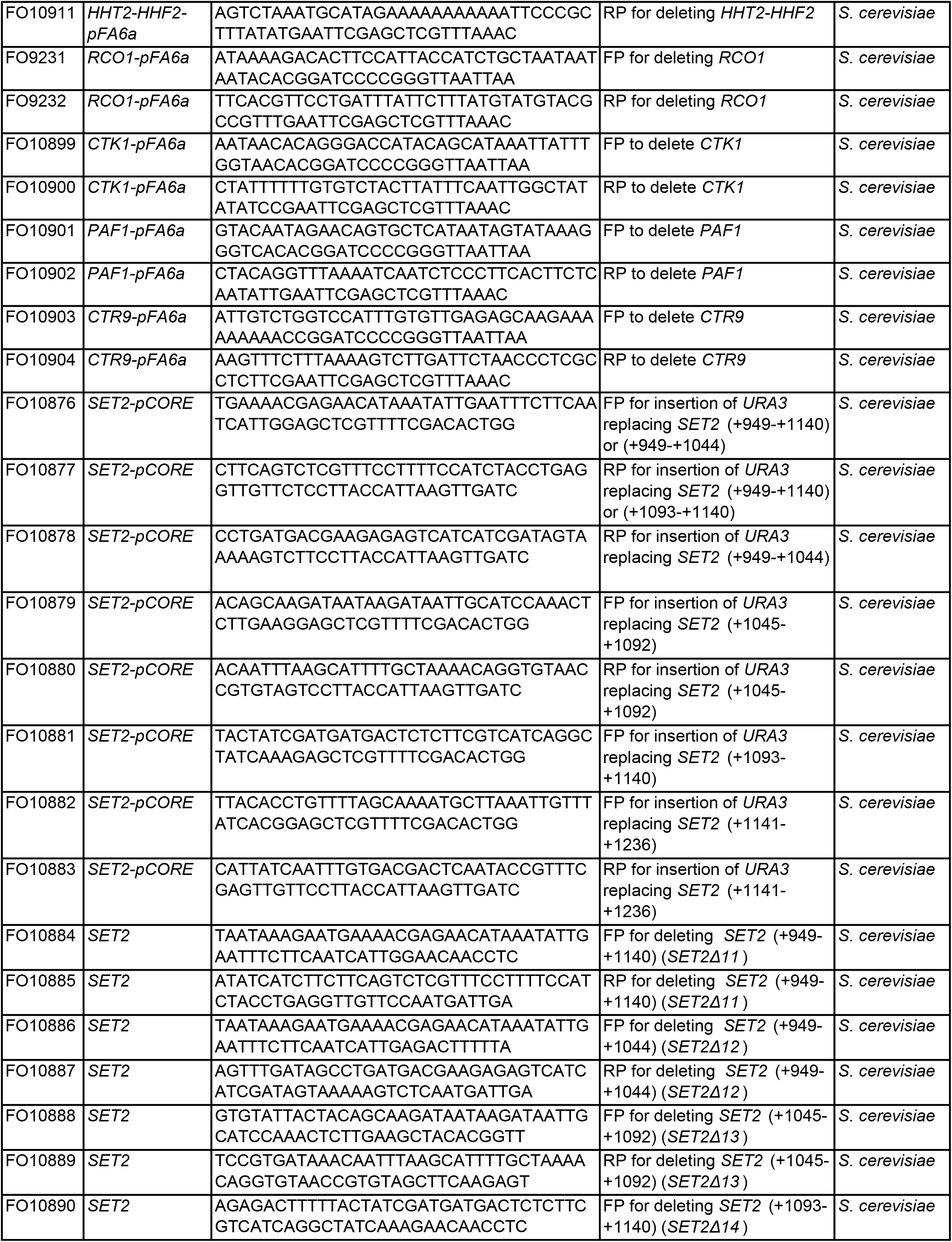
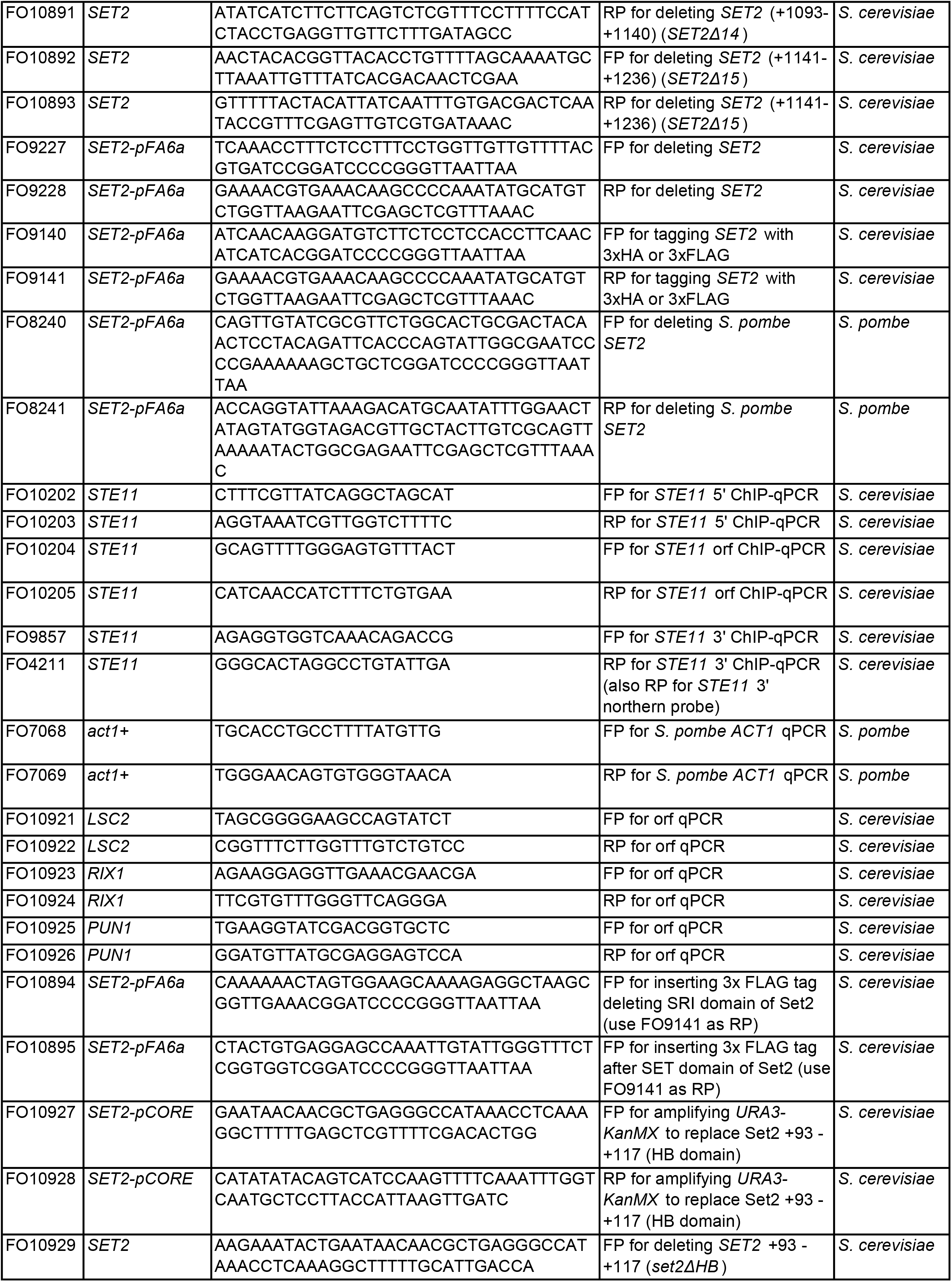
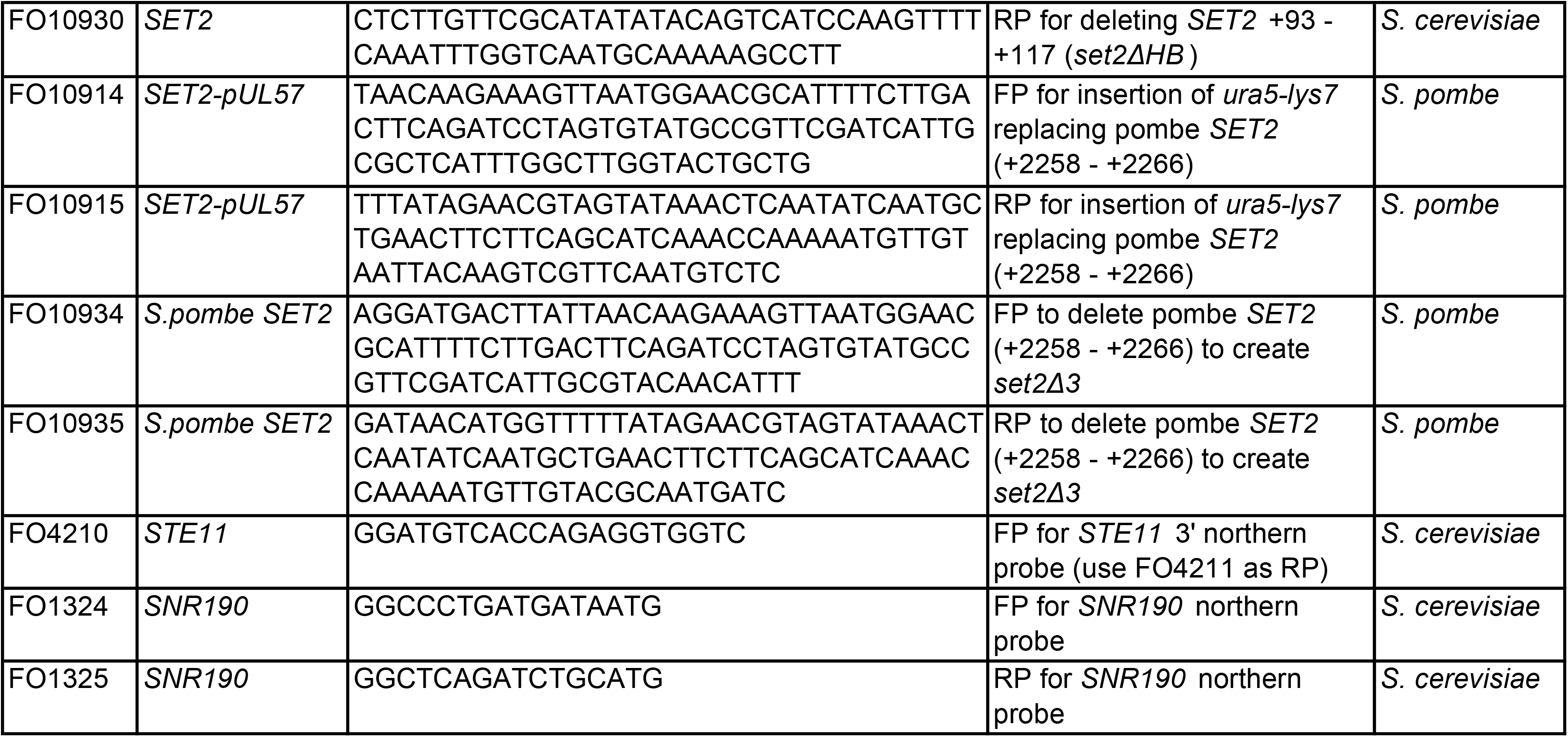

